# Reduced proteasome activity in the aging brain results in ribosome stoichiometry loss and aggregation

**DOI:** 10.1101/577478

**Authors:** Erika Kelmer Sacramento, Joanna M. Kirkpatrick, Mariateresa Mazzetto, Mario Baumgart, Aleksandar Bartolome, Simone Di Sanzo, Cinzia Caterino, Michele Sanguanini, Nikoletta Papaevgeniou, Maria Lefaki, Dorothee Childs, Sara Bagnoli, Eva Terzibasi Tozzini, Domenico Di Fraia, Natalie Romanov, Peter Sudmant, Wolfgang Huber, Niki Chondrogianni, Michele Vendruscolo, Alessandro Cellerino, Alessandro Ori

## Abstract

A progressive loss of protein homeostasis is characteristic of aging and a driver of neurodegeneration. To investigate this process quantitatively, we characterized proteome dynamics during brain aging in the short-lived vertebrate *Nothobranchius furzeri* combining transcriptomics and proteomics. We detected a progressive reduction in the correlation between protein and mRNA, mainly due to post-transcriptional mechanisms that account for over 40% of the age-regulated proteins. These changes cause a progressive loss of stoichiometry in several protein complexes, including ribosomes, which show impaired assembly / dis-assembly and are enriched in protein aggregates in old brains. Mechanistically, we show that reduction of proteasome activity is an early event during brain aging and is sufficient to induce proteomic signatures of aging and loss of stoichiometry *in vivo*. Using longitudinal transcriptomic data, we show that the magnitude of early life decline in proteasome levels is the major risk factor for mortality. Our work defines causative events in the aging process that can be targeted to prevent loss of protein homeostasis and delay the onset of age-related neurodegeneration.

**Highlights:**

- Progressive loss of stoichiometry affects multiple protein complexes
- Ribosomes aggregate in old brains
- Partial reduction of proteasome activity is sufficient to induce loss of stoichiometry
- Reduced proteasome levels are a major risk factor for early death in killifish

## Introduction

Although age is the primary risk factor for cognitive decline and dementia (Assoc, 2018), the associated age-dependent molecular changes are still not known in detail. Despite the presence of clear functional impairments (Buckner, 2004), physiological brain aging is characterized by limited loss of neurons (Schmitz and Hof, 2007) and specific morphological changes of synaptic contacts (Dickstein et al., 2013). Large collections of data for transcript dynamics in human and animal brains indicate that systematic, age-dependent changes in gene expression are also relatively minor (Cellerino and Ori, 2017), although some shared transcriptional signatures have been identified, including a chronic activation of cellular inflammatory response (Aramillo Irizar et al., 2018), reactive changes in glial cells (Clarke et al., 2018) and reduced expression of neuronal and synaptic genes (Lu et al., 2004; Somel et al., 2010).

Since the vast majority of human neurons are generated during fetal and perinatal life and neuronal turnover is limited in the postnatal human brain (Sorrells et al., 2018), neurons are particularly prone to accumulate misfolded proteins that are not properly processed by the cellular proteolytic mechanisms (proteasomal and autophagic pathways), thus forming aberrant deposits. Indeed, neurodegenerative diseases are characterized by the prominent presence of protein aggregates, in particular due to mutations that facilitate misfolding and aggregation, and impairment of cellular quality control systems (Soto and Pritzkow, 2018). Accumulation of protein aggregates occurs also during physiological aging, as demonstrated by the presence of lipofuscin (Glees and Hasan, 1976) and ubiquitinated cellular inclusions (Matsui et al., 2019; Zeier et al., 2011). However, the exact composition of these spontaneous aggregates and the mechanisms triggering their formation during brain aging remain unknown.

Although age-dependent transcript changes in the brain have been studied extensively (Blalock et al., 2003; Colantuoni et al., 2011; Loerch et al., 2008; Lu et al., 2004; Wood et al., 2013), we are just beginning to understand the corresponding global regulation of the proteome during aging (Ori et al., 2015; Somel et al., 2010; Walther et al., 2015). Substantial post-transcriptional regulation takes place in the aging brain, with a sizeable proportion of proteins being up or downregulated in the absence of changes in the levels of the corresponding transcripts (Ori et al., 2015), resulting a progressive mRNA-protein decoupling (Janssens et al., 2015; Wei et al., 2015). Protein aggregation could play a role in generating an imbalance between protein and transcript levels, but these aspects have not yet been investigated systematically in vertebrate brains.

To address this challenge, we studied the annual killifish *Nothobranchius furzeri*, which is the shortest-lived vertebrate that can currently be bred in captivity. With a lifespan of 3-7 months (Hu and Brunet, 2018; Ripa et al., 2017; Terzibasi et al., 2008; Valdesalici and Cellerino, 2003), it has emerged as a convenient model organism to investigate genetic and non-genetic interventions on aging (Cellerino et al., 2016; Harel et al., 2015; Kim et al., 2016; Platzer and Englert, 2016; Ripa et al., 2017), since it replicates many typical aspects of vertebrate brain aging at the levels of behaviour (Valenzano et al., 2006a; Valenzano et al., 2006b), neuroanatomy (Tozzini et al., 2012) and global gene expression (Aramillo Irizar et al., 2018; Baumgart et al., 2014). Age-dependent processes are enhanced in this species, thus facilitating the detection of differentially expressed genes as compared to other model organisms (Baumgart et al., 2014; Frahm et al., 2017; Wood et al., 2013). Importantly, an age-dependent formation of inclusion bodies containing α-synuclein and spontaneous degeneration of dopaminergic neurons has been recently described in killifish (Matsui et al., 2019). This phenotype closely mimics human pathologies and make killifish an extremely attractive vertebrate system to study age-related neurodegenerative disorders and therapeutic strategy against them.

In this work, we applied RNAseq, mass spectrometry-based proteomics and analysis of protein aggregates in killifish of different ages to delineate a timeline of molecular events responsible for loss of proteome homeostasis during brain aging. In particular, we set to identify the nature and biophysical properties of proteins that preferentially aggregate in old brains, to comprehensively investigate the loss of stoichiometry of protein complexes and their assembly state, and the role played by the proteasome as an early driver of protein homeostasis collapse using *in vivo* pharmacological experiments. Finally, we tested whether inter-individual differences in proteasome decline influence mortality.

## Results and Discussion

### Transcript and protein levels become progressively decoupled during brain aging

We initially analysed whole brains from animals of 3 different age groups by liquid chromatography tandem mass spectrometry using a label-free method. Based on previous phenotypic data, we chose to compare young, sexually mature fish (5 weeks post-hatching, wph), adult fish (12 wph) that do not show aging phenotypes (Terzibasi et al., 2008), and old fish (39 wph) that display neurodegeneration (Di Cicco et al., 2011; Terzibasi Tozzini et al., 2012) (Figure 1A and Table S1). Principal component analysis separated samples according to the age groups (Figure 1B). In order to achieve higher proteome coverage, we split the age groups into two separate experiments based on tandem mass tag (TMT) multiplexing, where we compared adult vs. young fish and old vs. adult fish (Figure S1A). This was necessary because of the limited number of channels available (10 per experiment) and to do not reduce the number of animals analysed per age group. A total of 8885 protein groups were quantified with at least two proteotypic peptides, of which 7200 were quantified in both experiments (Figure S1B). Almost half of the quantified protein groups (4179/8885) was significantly affected by aging in at least one of the age comparisons (Table S2). Functionally related proteins showed different patterns of abundance change between age groups, and pathways affected by aging in other species, including inflammation-related pathways (Aramillo Irizar et al., 2018), the complement and coagulation cascade (Clarke et al., 2018) were affected in killifish already in the transition from young to adult (Figure S1C and Table S3).

**Figure 1.**
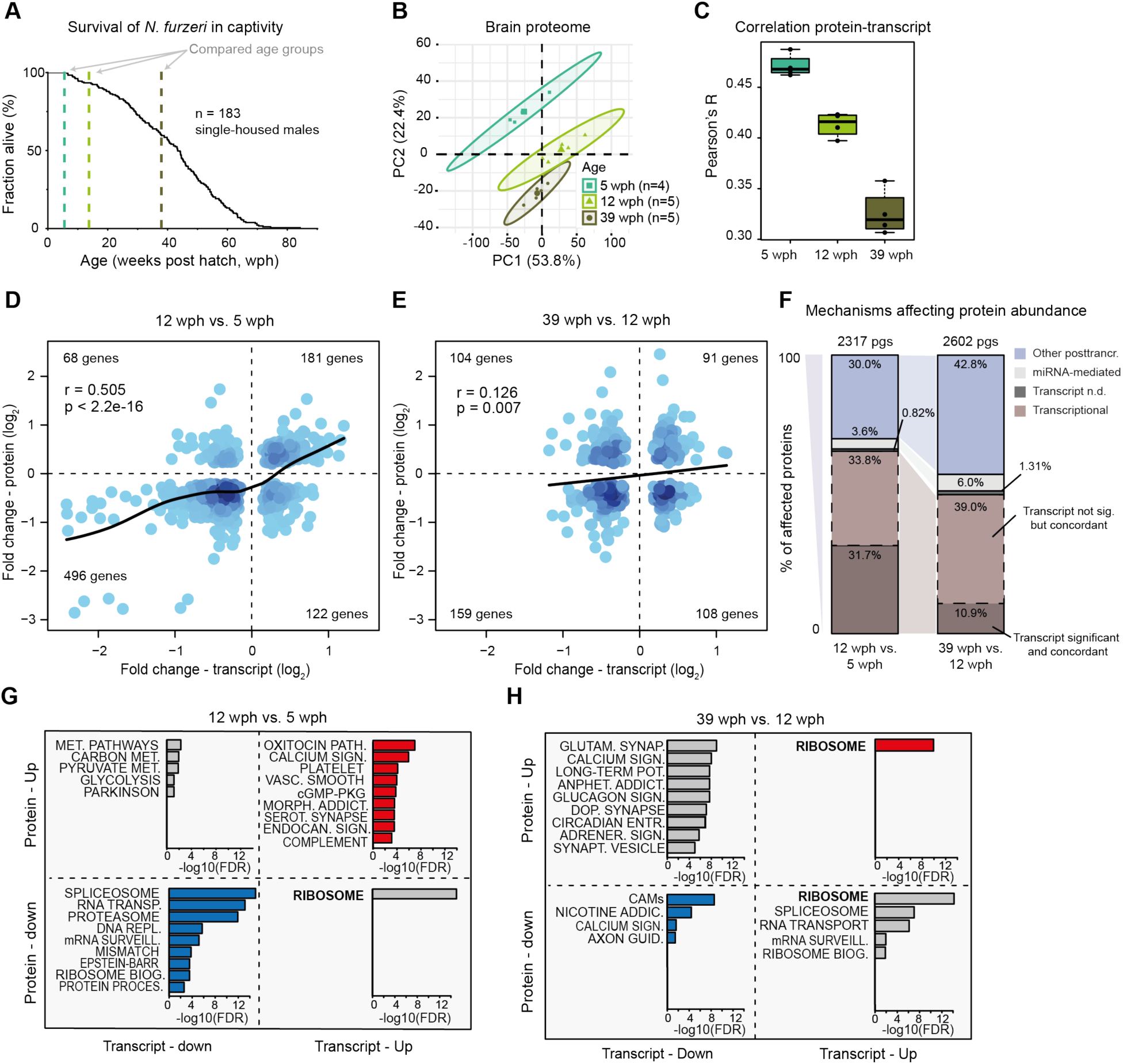
Transcript and protein levels become decoupled during *N. furzeri* brain aging. **(A)** Survival curve of *N. furzeri* in the FLI facility. Recording of deaths starts at age of 5 wph, which corresponds to sexual maturity, and the colored dashed lines indicate the three age groups analysed in this study (5 animals/group), namely 5 weeks post hatching (wph, young, sexual maturity), 12 wph (adult) and 39 wph (old, past median lifespan) of a wild-derived strain that exhibits a median lifespan of 7-8 months. **(B)** Principal component analysis (PCA) of brain samples based on the abundance of all proteins identified by label-free mass spectrometry. The smaller dots represent individual samples and the larger dots the centroids of each age-matched group. Ellipses represent 95% confidence intervals. The percentage of variance explained by the first two PC axes is reported in the axis titles. **(C)** Global protein-transcript correlation for each sample, grouped by age. RPKM and iBAQ values were used to estimate transcript and protein levels from matched RNAseq and TMT-based proteomics data obtained from the same animal. An ANOVA test was performed to evaluate significance among the age groups (mean correlation at 5 wph: 0.48; at 12 wph: 0.43; and at 39 wph: 0.33; p = 3.05e-07). **(D and E)** Scatter plot of log_2_ fold changes for genes differentially expressed both at transcript and protein levels (adj. p < 0.05). The color gradients indicate gene density in the regions where individual points overlap. Numbers of genes in each quadrant and the value of Pearson’s coefficient of correlation, r, are reported for each graph. Solid lines represent a spline fit (r = 0.505 for genes significantly affected at both transcript and protein levels, p < 2.2 × 10^−16^, D) (r = 0.126, p = 0.007, E). **(F)** Mechanisms affecting protein abundance during aging. Bar plots are based on all the proteins affected in either one of the age comparisons (adj. p < 0.05). Proteins were divided in the following five groups: (i) proteins and transcripts with significant and consistent changes (dark brown), (ii) proteins with significant changes, and with consistent changes of the transcripts (light brown), (iii) proteins with no transcripts detected (dark gray), (iv) proteins with transcripts whose translation is potentially regulated by miRNAs (light gray), as assessed by the workflow displayed in (Figure S2D), (v) all the remaining proteins that we classified as regulated by other post-transcriptional mechanisms (violet). pgs = protein groups. **(G and H)** Bar plots representing enriched KEGG pathways among genes that showed significant changes at both transcript and protein levels in aging. Genes were grouped according to the four possible patterns of transcript and protein regulation, as visualized by their positions in the four quadrants shown in (D) and (E), respectively. Only pathways significantly enriched (FDR < 0.05) are shown. The complete list of enriched pathways is reported in Table S5. Related to Figures S1 and S2, and Tables S2:S5.

Total RNAseq after rRNA depletion and microRNAseq were obtained from the same samples (Figure S1D and Table S4). For each sample, absolute protein abundances estimated from peptide intensities (iBAQ values, (Schwanhäusser et al., 2011)) were correlated with the corresponding transcript levels obtained by RNAseq (RPKM values), obtaining global protein-transcript correlation values for each sample separately. We observed a progressive age-dependent reduction of protein-transcript correlation values (Figure 1C), consistent with a decoupling between RNA transcripts and proteins during brain aging (Wei et al., 2015). Decoupling was observed also when analysing an independent RNAseq dataset from polyA+ RNA for animals of the same age groups (Baumgart et al., 2014) (Figure S1E). Fold-changes of genes differentially expressed in the two RNAseq datasets were strongly correlated (Figure S1F). For further analysis, we then focused on the dataset with higher sequencing depth and larger number of replicates, for which the absolute number of differentially expressed genes was higher (polyA+ RNA dataset, Figure S1G).

Direct comparison of protein and mRNA fold changes across age groups (Table S5) revealed discrepancies between RNA and protein regulation (Figure 1D, 1E and Table S2). Protein and transcript changes were significantly correlated in the adult vs. young fish comparison (Figure 1D), but the correlation was reduced in the old vs. adult comparison (Figure 1E), further supporting a progressive decoupling between transcript and protein regulation. For validation, we analyzed proteins previously identified to be very long-lived in rodent brain (Toyama et al., 2013), including histones, collagens and myelin proteins. For these proteins, we found that transcript, but not protein levels, were generally decreased in old fish, indicating that protein stability might contribute to the observed discrepancies between transcripts and proteins (Figure S2A). In contrast, protein levels estimated by mass spectrometry and immunoreactivity for the glial fibrillary acidic protein (GFAP) were shown to increase significantly in the aging brain (Terzibasi Tozzini et al., 2012), while RNA levels remained unchanged (Figure S2B). To exclude biases deriving from changes in cellular composition of the brain with aging, we analyzed the regulation of established cell markers (Sharma et al., 2015) in the age-group comparisons. We found only minor changes that were consistent at the protein and transcript level (Figure S2C), thus excluding that the observed decoupling between transcript and protein levels is due to changes in cellular composition.

To find out whether microRNAs could contribute to transcript-protein decoupling, we first analysed miRNA expression levels across the same three age groups (Table S4), then mapped the targets of age-affected miRNAs to our proteome data (Figure S2D). By considering potential regulation mediated by miRNAs, we defined a subset of proteins whose abundance is affected by aging via mechanisms independent of both transcript level and miRNA-mediated post transcriptional regulation (Figure 1F). This subset accounted for 30% of the affected proteins in the adult vs. young fish comparison, and increased up to 43% in the old vs. adult comparison.

To clarify whether the transcript-protein decoupling preferentially affects some specific pathways, we classified age-affected genes according to their respective transcript and protein fold changes, and performed pathway overrepresentation analysis (Table S5). In the comparison adult vs. young fish, pathways related to the complement coagulation cascade and synaptic function/plasticity were overrepresented in concordantly increased transcripts and proteins (Figure 1G), in agreement with the notion that synaptogenesis continues during this phase of residual brain growth (Tozzini et al., 2012). By contrast, genes coding for biosynthetic pathways such as RNA transport, splicing and surveillance of RNA, ribosome biogenesis, and protein processing in the endoplasmic reticulum (ER) were overrepresented in concordantly decreased proteins/transcripts (Figure 1G). These changes may be related to the reduction of adult neurogenesis (Tozzini et al., 2012) that accounts for a significant fraction of global transcription regulation occurring during this period (Baumgart et al., 2014). The same biosynthetic pathways become discordant when old and adult animals are compared, with protein levels decreasing further with age, while transcript levels changed directionality and increased (Figure 1H, bottom right quadrant).

Taken together, our data indicate that post-transcriptional mechanisms regulating protein levels have an increasingly important role in modulating protein abundance with age; they are responsible for nearly half of the protein changes observed in old brains, and they can lead to the opposite regulation trends for proteins and mRNAs.

### Loss of ribosome stoichiometry in the aging brain

Amongst genes showing opposite transcript and protein changes already in the adult vs. young fish comparison, we identified 13 genes encoding ribosomal proteins with transcript levels being significantly increased and protein abundances decreased (Figure 2A and Figure S3A). Fold changes of genes encoding for ribosomal proteins split into two groups in the old vs. adult fish comparison (Figure 1H): while transcript levels continue to increase consistently, ribosomal proteins show either increased (13 proteins, e.g., RPS20, RPL8 and RPL21) or decreased (14 proteins, e.g., RPS6, RPLP2 and RPL22L1) abundance (Figure 2A and S3A). A similar pattern was observed also for the mitochondrial ribosome (Figure S3B). These findings indicate a loss of stoichiometry of ribosomal proteins (i.e., an imbalance in their relative levels) during aging, which is likely to impair ribosome assembly and to create a pool of orphan proteins at risk of aggregation. When mapped on the ribosome structure (Khatter et al., 2015), age-affected proteins form clusters of either consistently increased or decreased abundance (Figure 2B). Since transcript level changes are consistent, while ribosomal protein levels are not (Figure S3A and S3B), the loss of ribosome stoichiometry must result from an alteration of post-transcriptional mechanisms mediating protein homeostasis. We obtained ribosome footprint data from young and old killifish brains and showed consistently increased levels of transcript encoding for ribosomal proteins to be associated with ribosomes in old brains, as previously shown in rats (Ori et al., 2015) (Figure S3C). These data exclude changes in translation output as a cause for the observed loss of ribosome stoichiometry, and point to other mechanisms such as protein degradation and aggregation.

**Figure 2.**
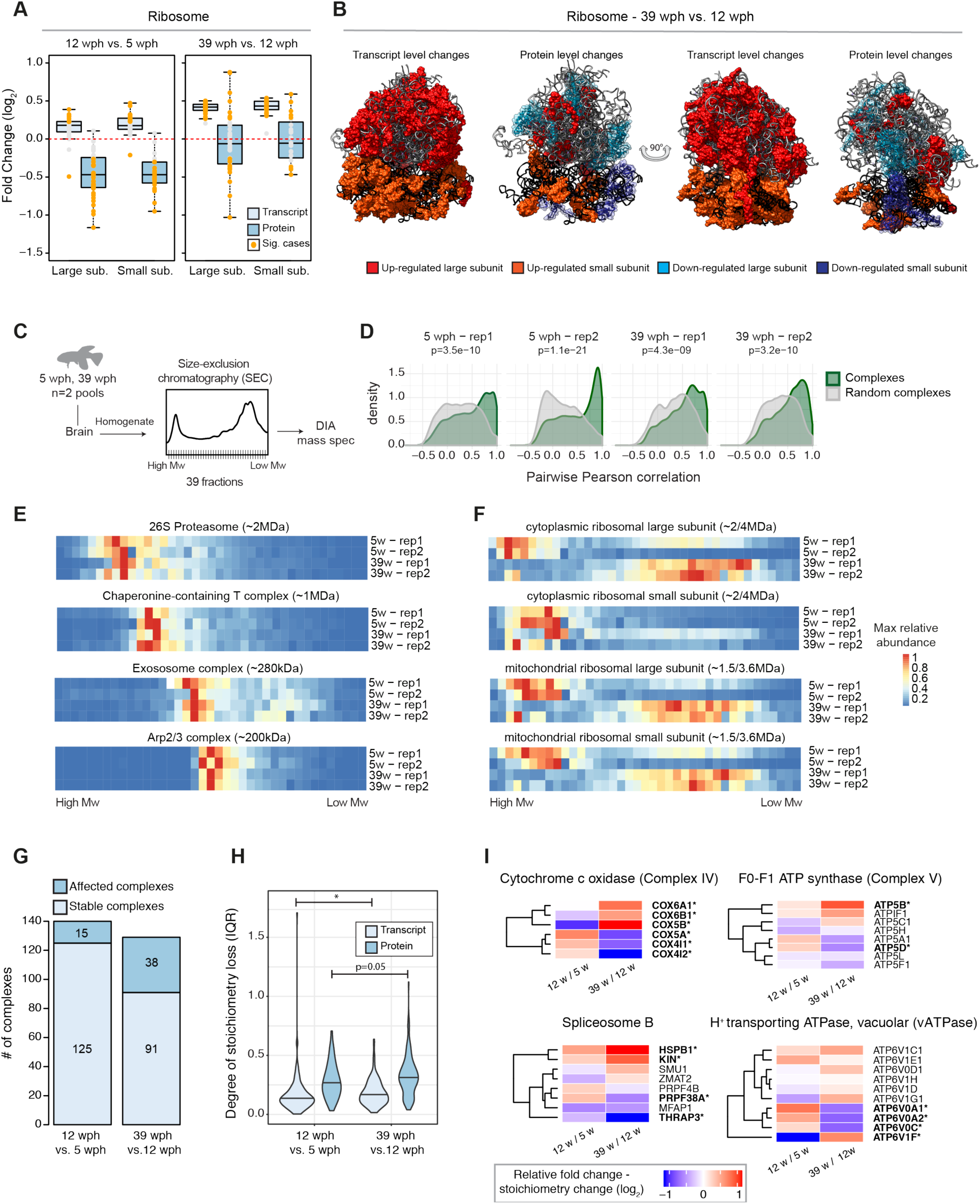
Loss of stoichiometry and disassembly of ribosomes in old killifish brain. **(A)** Abundance changes of ribosomal proteins and their transcripts during aging. Cytoplasmic ribosomal proteins of large and small subunits are displayed separately and changes are shown for both the age comparisons as box plots. Transcripts are displayed as light blue and proteins as dark blue boxes. Changes of individual proteins are displayed as dots; orange dots identify significant cases (adj. p < 0.05). **(B)** Visualization of age-related changes of proteins and transcripts projected on the 80S ribosome complex structure. Ribosomal RNAs are depicted in ribbon form: 28S rRNA, 5S rRNA, 5.8S rRNA of large subunit are depicted in light gray and 18S rRNA of small subunit is depicted in black. Ribosomal proteins are depicted as molecular surfaces and shown only if significant changes in the level of corresponding mRNA or protein were detected. Affected proteins of large and small subunits are visualized in 2 different shades of red (up regulated), or blue (down regulated). For clarity, down regulated components are displayed as transparent molecular surfaces. Visualization was performed with USCF Chimera program (version 1.12), according to Protein Data Bank archive - human 80S ribosome 3D model: 4UG0. (**C**) Brains from young (5 wph) and old (39 wph) were homogenized and clarified lysates separated by size-exclusion chromatography (SEC). For each age group, two pools of brains were processed separately. For each experiment (4 in total), 39 fractions were collected along the chromatogram, digested into peptides and analysed by Data Independent Acquisition (DIA) quantitative mass spectrometry. (**D**) Co-elution of members of protein complexes in SEC. For each experiment, the distribution of pairwise correlations between members of the same protein complex was analysed (green). As expected, members of protein complexes tend to co-elute in all SEC experiments, as indicated by positive correlation values. A set of randomly defined protein complexes was used as control (gray). For all the experiments, the correlations of real complexes are significantly higher than random ones, p < 2.2e-16 Wilcoxon Rank Sum test. **(E and F)** Co-elution profiles for selected protein complexes. For each complex, the median abundance of all the quantified subunits was used to generate the complex profile across fractions (Table S6). All the complex profiles are scaled to the max value (set to 1) to make profiles comparable across experiments. The estimated molecular weight of the displayed complexes is indicated in brackets. **(G)** Statistics of protein complexes undergoing stoichiometry changes with aging. Only protein complexes that had at least 5 members quantified were considered for each comparison. Complexes were considered affected if at least two members showed significant stoichiometry change (adj. p < 0.05 and absolute log_2_ fold change > 0.5). The complete list of stoichiometry changes is available in Table S7. **(H)** Violin plots depicting interquartile ranges (IQRs) of individual members of protein complexes during aging. The IQR for each protein complex considered in **G** was calculated using transcript (total RNA dataset, light blue) or protein (dark blue) log_2_ fold changes between two age groups. * p < 0.05, Wilcoxon Rank Sum test. **(I)** Heat map showing relative protein fold changes for members of selected complexes affected by aging. Names of significantly affected members in the 39 wph vs. 12 wph comparison (adj. p < 0.05 and absolute log_2_ fold change > 0.5) are highlighted in bold with a star. Related to Figure S3 and Tables S6 and S7.

In order to directly investigate the consequences of stoichiometry loss on the assembly state of ribosomes, we performed size-exclusion chromatography of brain lysates coupled to Data Independent Acquisition (DIA) quantitative mass spectrometry on 2 pools each of young and old killifish (Figure 2C). Our analysis retrieved known protein complexes as distinct co-eluting peaks in both young and old brains (Figure 2D and 2E, Table S6). Interestingly, we found protein components of the ribosome to co-elute at lower than expected molecular weight in old brain lysate. This effect was particularly pronounced for the large cytoplasmic ribosome and the mitochondrial ribosome (Figure 2F). Other complexes were not affected and eluted at the same retention time in both young and old lysates (Figure 2E), pointing to a specific effect on ribosomes. Taken together, these data indicate that age-dependent loss of stoichiometry of ribosomes might derive from altered assembly / dis-assembly in old brains.

### Widespread stoichiometric imbalance in protein complexes during aging

We next asked whether the age-related loss of stoichiometry described above occurs more widely in the proteome. We thus analysed all annotated protein complexes in the two age group comparisons (Ori et al., 2013; Ori et al., 2016). We found that the number of complexes undergoing stoichiometry changes increases from 11% (16 out of 140) between 5 and 12 wph, to 30% (39 out of 129) between 12 and 39 wph (Figure 2G and Table S7). Consistently, the number of affected complex members increases almost 2-fold in the old vs. adult comparison (from 6% to 13%) (Figure S3D). Loss of stoichiometry was confirmed by an alternative metric, namely an increase in the inter-quantile range (IQR) of fold changes of protein complex members (Janssens et al., 2015) in the old vs. adult fish comparison (Figure 2H). In order to exclude potential batch effects, we repeated the analysis on a subset of 3 animals for each age group that were analysed in the same TMT10plex experiment, and confirmed an age-dependent increase of IQR between protein complex members (Figure S3E).

When individual complexes were ranked according to the difference of IQR between the two age comparisons, the majority of the complexes showed an increase in IQR (76 out of 124, 61%, Figure S3F and Table S7). The most affected complexes included Complex IV and Complex V, but not Complex I and Complex III, of the mitochondrial respiratory chain, the cytoplasmic ribosome, the 26S proteasome, the B complex of the spliceosome, and the lysosomal V-type ATPase (Figure 2I). These complexes take part in biological processes known to be causative in aging (Carmona-Gutierrez et al., 2016; Chondrogianni et al., 2014; Dillin et al., 2002; Heintz et al., 2017; Lee et al., 2003; Steffen and Dillin, 2016), and the regulation of transcripts coding for them are correlated with individual lifespan in a longitudinal RNA-seq study in *N. furzeri* (Baumgart et al., 2016). Many of these complexes are affected by stoichiometry changes already at the adult stage (Table S7), identifying these alterations as early events during aging progression.

### Age-dependent protein aggregates are enriched for ribosomal proteins

Loss of stoichiometry and altered assembly of protein complexes can create a pool of orphan proteins at risk of aggregation. Since protein aggregates are known to be SDS-insoluble (Reis-Rodrigues et al., 2012), we compared SDS-insoluble fractions from brain homogenates of young and old animals (Figure 3A and Figure S4A). We used mice for this analysis because of the larger brain size that allows retrieval of sufficient amount of aggregates (that constitute only about 0.5% of total proteins in old brains) for proteomic analysis. As expected, the yield of SDS-insoluble protein aggregates was significantly higher from old animals, confirming that aging is associated with enhanced protein aggregation (Figure 3B and Figure S4B). We then analysed the aggregate composition by quantitative mass spectrometry to identify proteins enriched in these aggregates as compared to the starting total brain homogenates (Table S8). Enriched proteins showed a predicted higher molecular chaperone requirement for folding, and were richer in intrinsically disordered regions (Figure 3C). Among the enriched proteins, we found collagens (Col1a1, Col1a2 and Col4a2), which are well-known to undergo age-dependent crosslinking (Viidik, 1979), and ferritins (Fth1 and Ftl1), whose aggregation is linked to the age-dependent brain accumulation of intracellular iron (Ripa et al., 2017)(Figure 3D). Interestingly, protein aggregates were also enriched for ribosomal proteins (p=4.4e-05, Fisher test, Figure 3D and S4D) that are both characterised by protein transcript decoupling (Figure 1H) and loss of complex stoichiometry (Figure 2A and 2B) during aging. Other protein complexes that displayed loss of stoichiometry (i.e., Complex V and vATPase) did not show significant enrichment in aggregates, indicating that stoichiometry imbalance does not always correlate with protein aggregation (Figure S4D).

**Figure 3.**
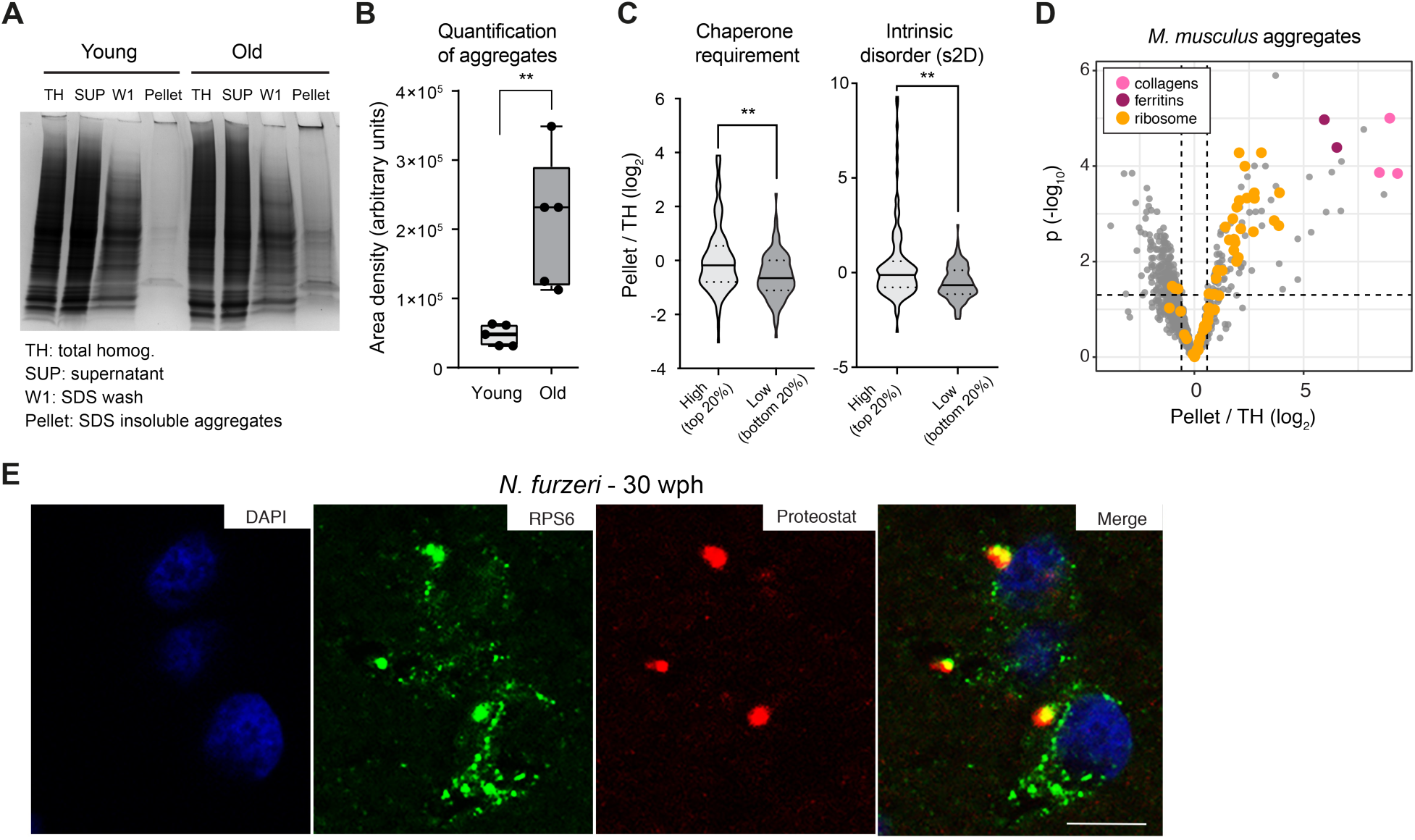
Aggregation of ribosomal proteins during brain aging. **(A)** Representative Coomassie-stained SDS-PAGE gel showing the isolation of SDS insoluble aggregates from mouse brain lysates. TH = total homogenate, SUP = supernatant, W1 = SDS-soluble fraction, Pellet = formic-acid soluble fraction (see Figure S4A). **(B)** Quantification of the yield of SDS-insoluble aggregates from young and old brain lysates was based on densitometry analysis of Coomassie-stained gel bands obtained from different animals, n=5 per age group (Figure S4B); ** p < 0.01, unpaired t-test. **(C)** Proteins enriched in aggregates show a predicted higher molecular chaperone requirement for folding (top vs. bottom 20% p=0.0055, Kolmogorov-Smirnov test), and are richer in intrinsically disordered regions (s2D-derived scores, top vs. bottom 20% p=0.0019, Kolmogorov-Smirnov test). Violin plots: the solid line shows the median, the dotted lines the interquartile ranges. The same result was obtained with cleverSuite-derived scores (top vs. bottom 20% p=0.0201, Kolmogorov-Smirnov test) (Figure S4C). **(D)** Volcano plot based on protein quantification by label-free mass spectrometry depicting the enrichment of specific proteins in protein aggregates. The x-axis indicates the log_2_ ratio between protein abundance in aggregates (Pellet) and starting total homogenate (TH). The horizontal dashed line indicates a p value cut-off of 0.05 and vertical lines a log_2_ fold change cut off of ± 0.5. Selected proteins are highlighted as colored dots as indicated in the figure legend. Protein quantification was based on samples obtained from 3 independent isolations. **(E)** Double-labelling of telencephalic sections of *N. furzeri* with anti-RPS6 (green) as ribosomal marker and Proteostat as a marker for aggregated proteins (red). Nuclear counterstaining was performed with DAPI (blue). Scale bar = 10 µm. Related to Figure S4 and Table S8.

To confirm the aggregation of ribosomal proteins in killifish, we performed staining of young (7-10 wph) and old (27-30 wph) brain slices using Proteostat, an amyloid-specific dye (Shen et al., 2011). As expected, we detected lysosomal aggregates in old, but not in young brains (Figure S4E, S4F and S4G). These aggregates appeared to contain the ribosomal protein RPS6 (Figure 3E and S4H), which was found to be significantly enriched in aggregates in mice (Table S8). In addition, we performed mass spectrometry on aggregates from killifish. Although the limited amount of material precluded a quantitative analysis as performed in mouse, we were able to confidently identify several ribosomal proteins also in killifish brain aggregates (Figure S4I and Table S8). Taken together, these data demonstrate that aggregation of ribosomal proteins is a conserved trait of brain aging in fish and mice.

### Acute partial reduction of proteasome activity is sufficient to induce loss of protein stoichiometry in vivo

Protein stoichiometry loss could be due to decreased proteolysis rates. We focused on the proteasome, which is one of the main degradation machineries and itself undergoes stoichiometry loss upon aging in killifish (Figure S3F). First, we observed a decrease in the levels of both proteasome proteins and their transcripts in adult killifish, which was followed by a stoichiometry imbalance that manifested in old fish exclusively at the protein level. In particular, we observed an imbalance between proteins belonging to the 19S and the 20S complexes, with the latter being exclusively upregulated in old fish (Figure 4A and 4B). We confirmed an age-dependent increase of 20S relatively to single- (26S) and double- (30S) capped proteasomes using immunoblots based on native gel electrophoresis from an independent cohort of samples (Figure 4C and S5A).

**Figure 4.**
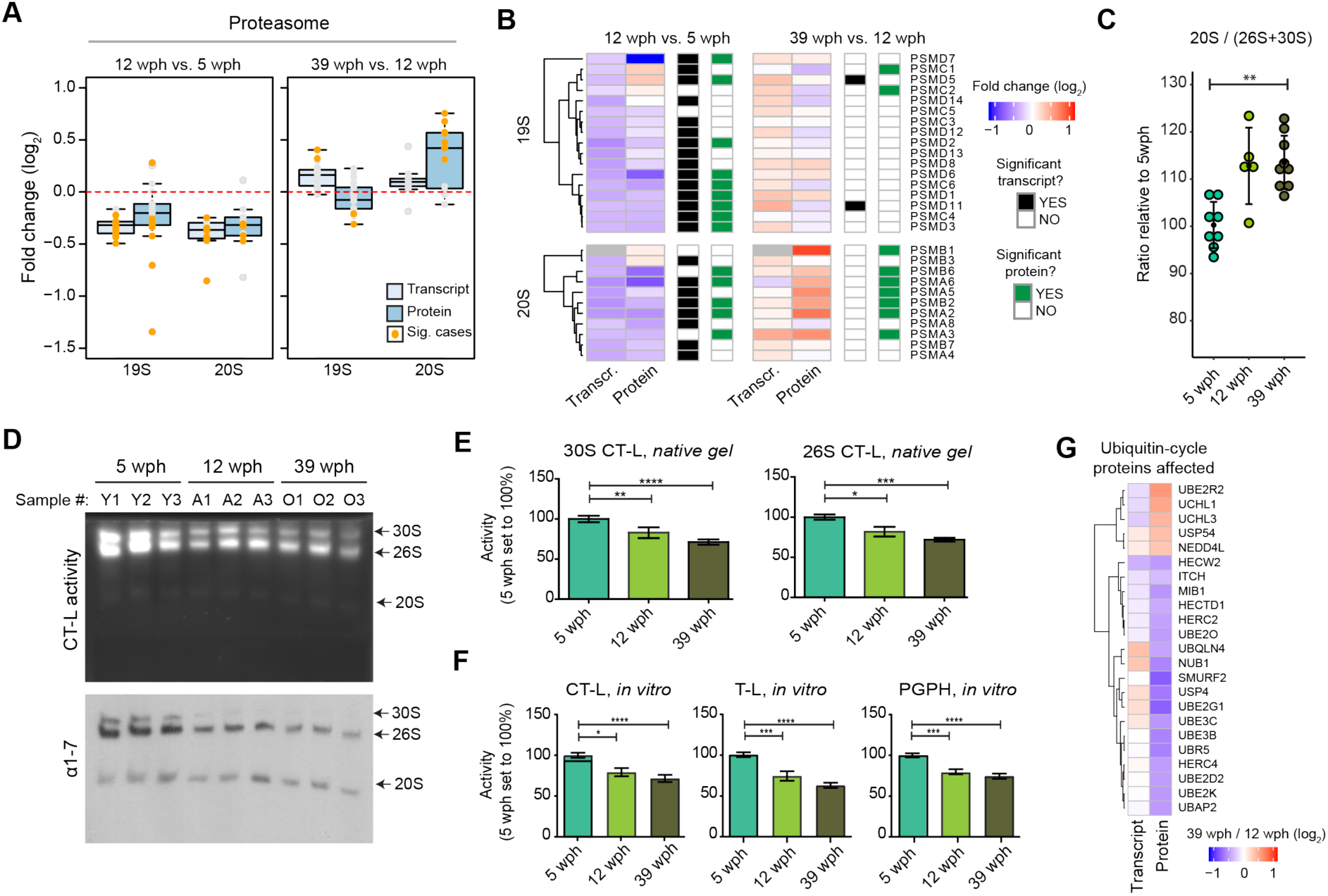
Reduced proteasome activity and assembly in old brains. **(A)** Abundance changes of proteasome proteins and their transcripts during aging. Members of the 19S and 20S complex are displayed separately and changes are shown for both the age comparisons as box plots. Transcripts are displayed as light blue and proteins as dark blue boxes. Changes of individual proteins are displayed as dots, orange dots represent significant cases (adj. p < 0.05). **(B)** Heat map showing transcript and protein fold changes for members of the 26 proteasome (19S and 12S complexes). Genes are annotated according to significance of their changes at the level of transcript (adj. p < 0.05: black, adj. p > 0.05: white) or protein (adj. p < 0.05: green, adj. p > 0.05: white). **(C)** The between 20S and 26S+30S proteasome abundance assessed by immuno blot on native gels (Figure S5A). ** p<0.001, Wilcoxon Rank Sum test. **(D)** In-gel proteasome assay following native gel electrophoresis (top) and immunoblotting of proteasome complexes (30S, 26S and 20S) (bottom) in young (5wph), adult (12 wph) and old (39 wph) killifish brains. For additional samples and low exposure pictures see Figure S5B and S5C. **(E)** Bar plots depicting the quantification of chymotrypsin-like (CT-L) activity from native gels calculated for doubly capped (30S) or singly capped (26S) proteasomes. n >=5 per sample group; error bars indicate standard deviation. * p < 0.05, ** p < 0.01, *** p < 0.001, unpaired t-test. For each sample group, the mean value of activity in young samples (5 wph) was set to 100%. **(F)** Percentage (%) of chymotrypsin-like (CT-L), trypsin-like (T-L) and peptidylglutamyl peptide hydrolysing or caspase-like (PGPH) proteasome activities in brain extracts of killifish of different ages. n >=6 per sample group; error bars indicate standard deviation. * p < 0.05, ** p < 0.01, *** p < 0.001, ANOVA. For each sample group, the mean value of each activity in young samples (5 wph) was set to 100%. **(G)** Age-related changes of proteins involved in the ubiquitin cycle. All the displayed proteins showed significant protein level changes in the 39 wph vs. 12 wph comparison (adj. p < 0.05). Related to Figure S5.

To further investigate the proteasome functional status in killifish brains of different age groups, we performed native gel electrophoresis of proteasomes accompanied by in-gel proteasome activity assays (Chondrogianni et al., 2015). A significant decrease in the levels of both 30S and 26S proteasomes was revealed already in adult animals (Figure 4D, S5B and S5C). This decrease was accompanied by a significant reduction of all three proteasome activities in adult and old samples as compared to young samples (Figure 4E and 4F). Additionally, we detected downregulation of enzymes involved in the ubiquitin cycle (ubiquitin-conjugating enzymes and ubiquitin ligases) in the old killifish brain (Figure 4G). These include the ubiquitin ligase UBE2O that has been shown to mediate recognition, ubiquitination and targeting for proteasomal degradation of mislocalized ribosomal proteins (Yanagitani et al., 2017).

Next, we investigate whether an acute partial reduction of proteasome activity is sufficient to induce age-relate phenotypes in killifish brain. Thus, we treated adult (12wph) killifish with Bortezomib, a reversible proteasome inhibitor, for four days and achieved ∼50% inhibition of proteasome activity in brain, mimicking the activity observed in old fish (Figure 5A and 5B). Quantitative mass spectrometry revealed distinct proteome changes induced by partial proteasome inhibition in brain (Figure 5C and Table S9). These included a compensatory up-regulation of proteasome activators (e.g., PSME4), ubiquitin ligases (e.g., UBE3C), autophagy-related proteins (e.g., ATG2B) and heat shock proteins (e.g., HSPB1) (Figure 5D). Interestingly, this proteomic signature mimicked an aging phenotype, as indicated by a significant overlap with proteins whose abundance changes with age (Figure 5E). Using the same approach applied for aging data, we were able to detect protein complexes whose stoichiometry was affected by acute proteasome inhibition. These include the large subunits of both cytosolic and mitochondrial ribosomes (Figure 5F, 5G and Table S9). Taken together, these data indicate that decreased proteasome activity is an early event during brain aging that is sufficient to induce loss of stoichiometry of ribosomes in the brain *in vivo*.

**Figure 5.**
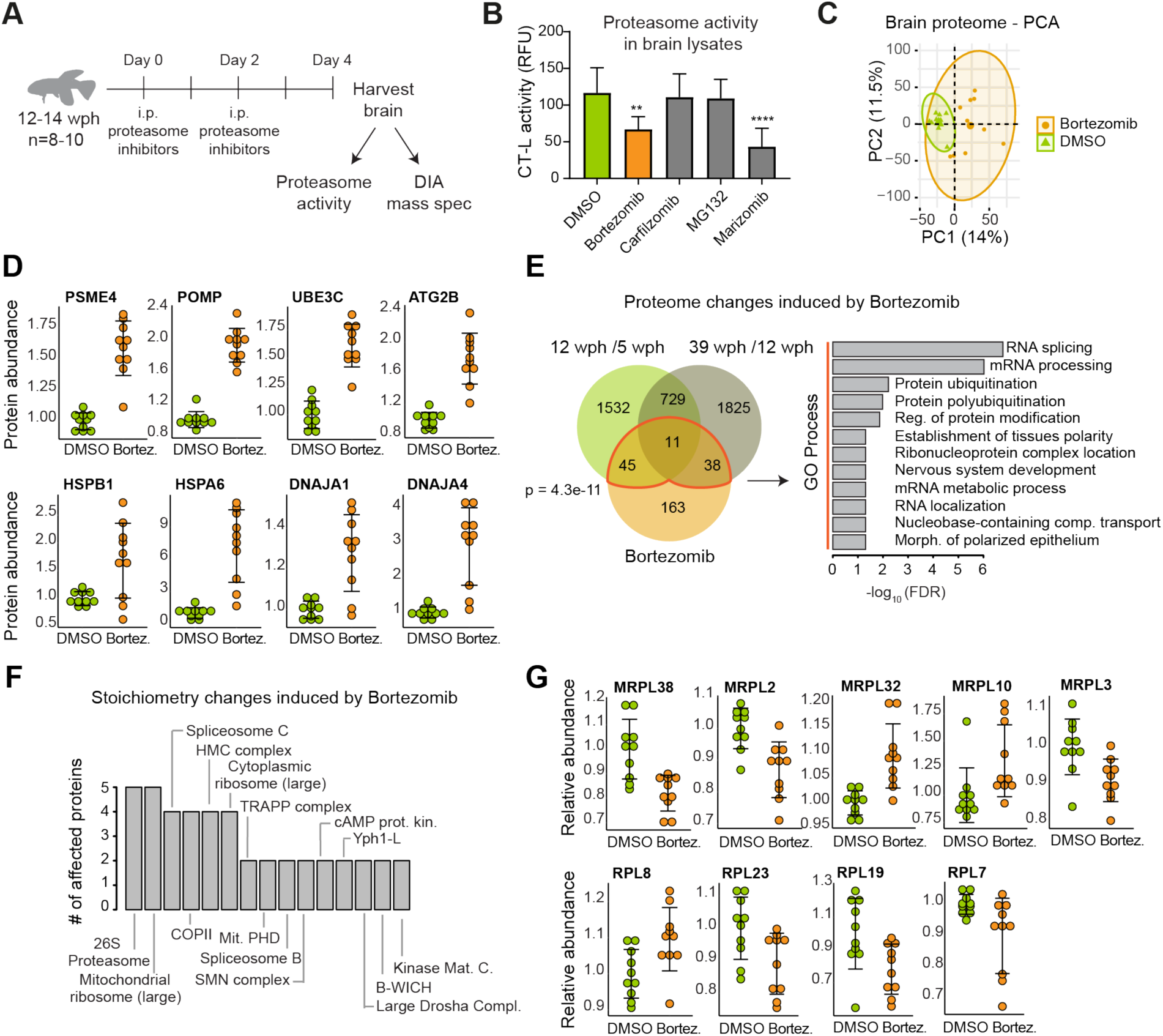
Partial pharmacological inhibition of proteasome activity in adult killifish brain affects aging protein networks and induces stoichiometry changes in a subset of protein complexes. (**A**) Study design. Adult (12-14 wph) killifish were treated with different proteasome inhibitors or vehicle control for 4 days. Proteasome activity and proteome changes were analysed in brains from treated and control fish. (**B**) Chymotrypsin-like (CT-L) proteasome activity in brain extracts of killifish treated with different proteasome inhibitors. The activity was measured at day 4 after the beginning of treatment. n=8 per sample group; error bars indicate standard deviation. ** p < 0.01, **** p < 0.0001, one-way ANOVA followed by Holm-Sidak’s multiple comparisons test. (**C**) Principal component analysis (PCA) of brain samples based on proteome profiles obtained by Data Independent Acquisition (DIA) quantification treated with Bortezomib or vehicle control (DMSO). n=10 per sample group. The smaller dots represent individual samples and the larger dots the centroids of each age-matched group. Ellipses represent 95% confidence intervals. The percentage of variance explained by the first two PC axes is reported in the axis titles. (**D**) Proteasome inhibition induces proteasome activators (PSME4), assembly factors (POMP) ubiquitin ligases (UBE3C), mediators of autophagosome formation (ATG2B), heat shock proteins and chaperones in killifish brain. Protein abundances were quantified by DIA mass spectrometry and they are shown relative to the mean value of vehicle control samples (DMSO) set to 1, n=10 per sample group. Adj. p < 0.05 for all the displayed proteins. (**E**) Overlap between proteins affected by Bortezomib treatment and aging in killifish brain. For all comparisons, only significantly affected proteins adj. p < 0.05 were considered. A significant overlap between Bortezomib and aging-affected proteins is detected (Fisher test, as indicated in the figure panel). Significantly enriched GO biological process terms in the subset of overlapping proteins are indicated (FDR < 0.05). (**F**) Bortezomib treatment affects the stoichiometry of a subset of protein complexes. The number of affected proteins (adj. p < 0.25) for each protein complex is indicated. Only protein complexes that had at least two members affected are shown. (**G**) Members of the mitochondrial and cytoplasmic ribosomes affected by Bortezomib treatment. Relative protein abundances (normalized to the mean of the protein complex to which they belong) are shown. The mean value of vehicle control samples (DMSO) was set to 1. n=10 per sample group. Adj. p < 0.25 for all the displayed proteins. Related to Table S9.

### Early-in-life decrease of proteasome is the major risk factor for early death

To assess whether decline of proteasome levels is relevant for lifespan determination, we analyzed a longitudinal dataset of RNA-seq comprising 159 killifish where transcripts from fin biopsies were quantified at 10 and 20 wph. Age-dependent variations in gene expression were related to the lifespan of individual fish (Figure 6A). By implementing a Cox-Hazard model, we identified genes for which the amplitude of age-dependent regulation significantly correlates with mortality risk (Figure 6B and Table S10). We found the proteasome to be the most enriched category among genes predictive of lifespan (Figure 6C). In particular, a decrease of expression of proteasomal transcript between 10 and 20 wph increases mortality risk, and is, thereby, associated with shorter lifespan. Accordingly, when we classified the 159 fish on the basis of changes in proteasome expression, we found that the lifespan of individuals showing the largest age-dependent downregulation of transcripts coding for proteasomal proteins was significantly shorter than the lifespan of individuals showing the largest up-regulation (Figure 6D), These data support the finding obtained in the brain that the decrease of proteasome levels is an early event during aging and demonstrate that the rate of proteasome downregulation in early adult life is predictive of lifespan in killifish.

**Figure 6.**
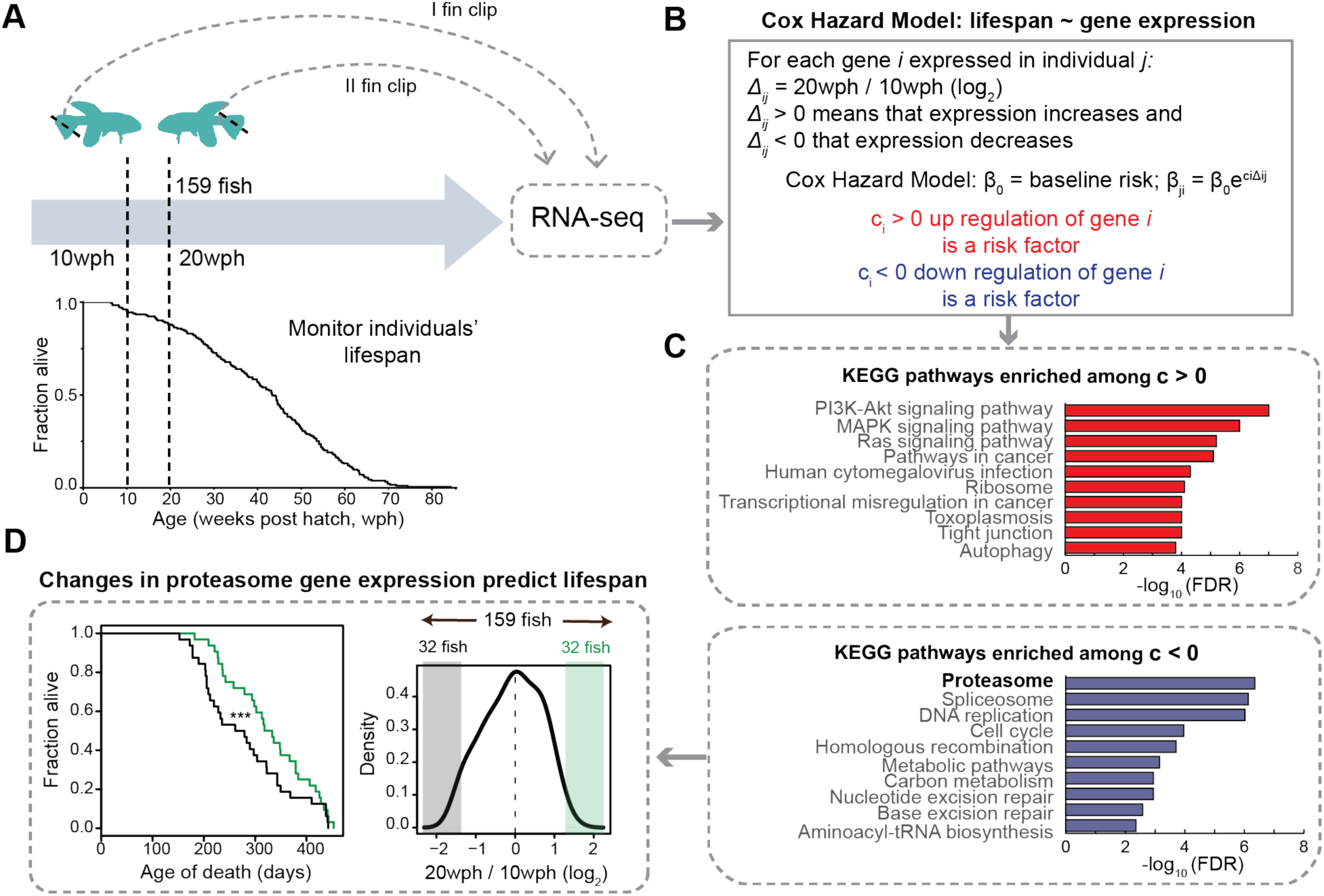
Longitudinal study in 159 killifish identifies early in life decrease of proteasome transcripts as a major risk factor for reduced lifespan. (**A**) Study design. Two fin clips were taken at 10 and 20 weeks post hatching (wph) from 159 killifish and individual lifespans monitored. RNA sequencing was employed to compare transcriptome changes between 10 and 20 wph for each individual fish. (**B**) A Cox Hazard model was used to correlate lifespan to gene expression changes. Two groups of genes were identified: (i) genes whose increased expression between 20 and 10 wph is a risk factors (i.e., associated to increased mortality risk) (red), and (ii) genes whose decreased expression is a risk factors (blue). (**C**) KEGG pathways enriched among genes whose regulation is associated to mortality. Only pathways with FDR<0.05 are shown. (**D**) Distribution of change in expression for proteasome transcripts across the entire cohort of 159 fish. Lifespan was compared among fish that showed extreme changes in proteasome levels between 10 and 20 wph (32 fish showing the most pronounced decreases, shown in black vs. 32 fish showing the most pronounce increases). *** p<0.001 Log-rank test. Related to Table S10.

### A molecular timeline for aging

Our results delineate a timeline of events associated with loss of protein quality control during aging. An early event, detectable already in adult fish, is a decreased proteolytic activity of the proteasome, which is driven by a downregulation of transcripts coding for components of the 19S and 20S complexes in adult fish brain (Figure 4A and 4B). The amplitude of this downregulation correlated with individual lifespan and could represent an early driver of aging (Figure 6D). The reduction of proteasome activity precedes chronologically the decoupling of transcript/protein levels, suggesting a causative role in this aspect of the aging process. Decreased proteasome activity can lead to the accumulation of proteins that are synthesized in excess relative to their binding partners, thus causing a stoichiometric imbalance of protein complexes (McShane et al., 2016). For instance, deletion of the ubiquitin ligase Tom1 (yeast homologue of Huwe1), which is responsible for the labelling for degradation of overproduced ribosomal proteins, leads to accumulation of multiple ribosomal proteins in detergent-insoluble aggregates in yeast (Sung et al., 2016). Accumulation of ribosomes in detergent-insoluble aggregates (David et al., 2010) and a loss of stoichiometry in the proteasome (Walther et al., 2015) were previously reported to occur during aging in *C. elegans.* Our work demonstrates the conservation of these mechanisms in the vertebrate brain, by showing alteration of stoichiometry in several large protein complexes (Figure 2G-I) and aggregation of ribosomes in old brain (Figure 3D and 3E). Specifically, we establish a mechanistic link between the partial reduction of proteasome activity observed in adult brains and the loss of stoichiometry of protein complexes (Figure 5F and 5G).

Later in life, the stoichiometry imbalance in protein complexes contributes to exacerbate the loss of protein homeostasis. Proteasome activity is further reduced in the old brain, correlating with an increased imbalance between the 19S and 20S complexes with time (Figure 4B and 4C). In addition, altered stoichiometry of the ribosome can underlie both the reduction and the qualitative changes of protein synthesis in brain aging (Ori et al., 2015; Schimanski and Barnes, 2010; Sudmant et al., 2018). An alteration of the stoichiometry between membrane-bound and cytosolic components of the lysosomal v-type ATPase (Figure 2I) might influence the acidification of lysosomes and the activation of mTORC1 (Zoncu et al., 2011), thus hampering the clearance of protein aggregates. These aggregates in turn may further impair the proteasome activity (Grune et al., 2004), thus creating a negative feedback loop. The interconnectivity between proteasome and lysosome/autophagy system is further highlighted by the fact that partial inhibition of the proteasome in the adult brain induces mediators of autophagosome formation (e.g., ATG2B) and autophagy receptors (e.g., SQTSM/p62), likely as a compensatory mechanism to ensure removal of non-degraded proteins at risk of aggregation. It is tempting to speculate that a progressive decline of proteasome activity might be the trigger for the impairment of lysosomal function that characterize aging and late-onset neurodegenerative disorders (Wallings et al., 2019). The combination of reduced proteasome activity and impaired lysosome/autophagy would make old brains more vulnerable to the accumulation of protein aggregates, neuronal loss and, consequently, favor the onset of neurodegenerative disorders.

Other key pathways implicated in aging are affected by loss of stoichiometry: in particular, alterations of respiratory chain complexes (particularly Complex IV and V) might contribute to their decreased activity and increased ROS production in old brain (Stefanatos and Sanz, 2018), and changes in multiple spliceosome complexes might underlie previously observed qualitative changes of splicing (Mazin et al., 2013; Ori et al., 2015). More detailed mechanistic studies are needed to demonstrate whether the alterations that we describe contribute to functional impairment of these protein complexes during aging or, rather, they represent adaptive responses to the aging process itself.

It remains to be determined which mechanisms promote the early decrease of proteasome activity in adult fish. Our data point to multiple processes being involved, including in particular: (i) decreased levels of rate-limiting proteasome members for the production of 20S assembled/functional proteasomes (e.g., PSMB5 or PSMB6, (Chondrogianni et al., 2015)), which we found to be significantly decreased already in the adult fish (Figure 4B); (ii) changes in abundance of proteasome proteins that are important for the assembly and activity of the 19S proteasome complex, such as PSMD5. PSMD5 has been also shown to inhibit the assembly and activity of the 26S proteasome, and this proteasome member has been shown to be induced by inflammation (Shim et al., 2012). Accordingly, we detected also in killifish an activation of inflammation-related pathways (Figure S1C) and, importantly, we identified PSMD5 as one of the few proteasome members to be upregulated in the adult fish (Figure 4B). Finally, PSMD11 (known as RPN-6 in *C. elegans*), which has been shown to be responsible for increased proteasome assembly and activity in human embryonic stem cells and in *C. elegans* (Vilchez et al., 2012), was significantly downregulated already in adult fish (Figure 4B). Identifying ways of counteracting these mechanisms might provide new avenues to delay organ dysfunction in aging and to increase lifespan. In this context, multiple studies have reported that transgenic animals (from various species) engineered to have enhanced proteasome activity show increased health- and lifespan (Augustin et al., 2018; Chondrogianni et al., 2015; Vilchez et al., 2012), and that proteasome activity is preserved in cells from centenarians (Chondrogianni et al., 2000). Correspondingly, we have shown that expression level of proteasome genes predicts life expectancy in killifish (Figure 6C and 6D).

In conclusion, our work identifies the maintenance of proteasome activity upon aging as being critical to ensure the correct stoichiometry of protein complexes involved in key biological functions such as protein synthesis, degradation, and energy production.

## Acknowledgments

The authors gratefully acknowledge support from the FLI Core Facilities Proteomics, Sequencing, Imaging and Bioinformatics, and the Fish Facility. The authors would like to acknowledge Stephan Schacke for assistance in data visualization, Toby Mathieson and Holger Dinkel for assistance in establishing data analysis pipelines, Matthias Platzer for support of longitudinal RNA-seq experiment, Sabine Matz and Bernhard Schlott for assistance with experiments, and Martin Beck, K. Lenhard Rudolph, David M. Sabatini, Maria Ermolaeva, Monther Abu-Ramaileh and Aliaksandr Khaminets for critical comments on the manuscript and helpful discussion. The FLI is a member of the Leibniz Association and is financially supported by the Federal Government of Germany and the State of Thuringia. AO and SDS acknowledge funding from the German Research Council (Deutsche Forschungsgemeinschaft, DFG) via the Research Training Group ProMoAge (GRK 2155). Research from NC lab is currently co-financed by the European Union and Greek national funds through the Operational Program Competitiveness, Entrepreneurship and Innovation under the call RESEARCH – CREATE – INNOVATE (project codes: T1EDK-00353 and T1EDK-01610) and under the Action “Action for the Strategic Development on the Research and Technological Sector” (project STHENOS-b, MIS 5002398). NP receives a PhD fellowship from Empirikion Foundation.

## Author contributions

Conceptualization: EKS, JMK, MM, NC, MV, AC, AO. Data curation: EKS, JMK, MM, AO. Formal analysis: EKS, JMK, MM, DC, MS, NR, MB, DDF, PS, NC, AO. Investigation: EKS, JMK, SDS, CC, NP, ML, ETT, NC. Methodology: EKS, JMK, MM, DC. Project administration: AC, AO. Resources: MB. Data analysis: MM, MS, DC, NR, PS, WH, AO. Supervision: JMK, WH, NC, MV, AC, AO. Visualization: EKS, JMK, MM, SDS, MS, DC, SB, ETT, AB, DDF, AO. Writing – original draft: EKS, JMK, MM, MV, AC, AO. Writing – review & editing: MS, ETT, NR, NC.

## Supplemental Information

Materials and Methods

Figures S1-S5

Table S2

Tables S1, S3-S10 (provided as separate Excel files)

Supplemental References

