## Supplementary Figures and Material and Methods for "Reduced proteasome activity in the aging brain results in ribosome stoichiometry loss and aggregation"

**This PDF file includes:**

Materials and Methods
Figures S1 to S5
Table S2
Captions for Tables S1, S3-S10 (provided as separate Excel files)

**Other Supplementary Materials for this manuscript include the following:**

Table S1: Experimental animals
Table S3: *N. furzeri* proteome aging
Table S4: *N. furzeri* transcriptome aging
Table S5: Comparison of transcriptome and proteome (*N. furzeri*)
Table S6: Size Exclusion Chromatography – mass spectrometry (*N. furzeri*)
Table S7: Protein complex analysis for *N. furzeri*
Table S8: Protein aggregates (*M. musculus* and *N. furzeri*)
Table S9: *In vivo* proteasome inhibition – proteome and protein complex analysis (*N. furzeri*)
Table S10: Longitudinal study (*N. furzeri*)

#### Materials and Methods

1. *Experimental animals including strain and housabandry details*

1.1. Fish maintenance

The *N. furzeri* strain was maintained at the FLI facility as described in (Baumgart et al., 2014). To avoid effects of circadian rhythms and feeding, animals were always sacrificed at 10 am in fasted state. For tissue preparation, fish were euthanized with MS-222 (PharmaQ) and cooled on crushed ice. Animals used for *in vivo* pharmacological interventions (section 3) were euthanized by rapid-chilling. A complete list of fish used for the different experiments is reported in Table S1.

1.2. *Mouse maintenance*

All mice were C57BL/6J obtained from Janvier or from internal breeding. Animals were maintained at the Leibniz Institute on Aging – Fritz Lipmann Institute (FLI) in a specific pathogen-free animal facility with a 12 h light/dark cycle. Mice were euthanized with $CO_2$. A complete list of mice used for the different experiments is reported in Table S1.

1.3. *Ethical Statements*

All experiments were performed in accordance with relevant guidelines and regulations. Fish were bred and kept in FLI´s fish facility according to §11 of German Animal Welfare Act under license number J-003798. The protocols of animal experimentations were approved by the local authority in the State of Thuringia (Veterinaer- und Lebensmittelueberwachungsamt; proteasome inhibition: reference number 22-2684-04-FLI-19-010). Sacrifice and organ harvesting of non-experimental animals was performed according to §4(3) of German Animal Welfare Act.

2. *Size exclusion chromatography (SEC) coupled to mass spectrometry*

2.1. *Sample preparation*

Individual brains of young (5 wph) and old (39wph) fish were collected and snap frozen in liquid nitrogen (Table S1). On the preparation for SEC, at least 4 brains were pooled for each replicate in order to obtain approx. 3 mg of protein extract as starting material, and lysed in 1.5 mL of lysis buffer (50 mM HEPES, pH 6.8, 1 mM MgCl2, 1 mM DTT, 150 mM NaCl, 5 mM ATP, cycloheximide 100 µg/ml, RNAse inhibitor 50U, protease and phosphatase inhibitors). Samples were then vortexed (5 times) prior to sonication using a Bioruptor Plus (Diagenode) for 5 cycles (30 sec ON/60 sec OFF) at high setting, at 4 °C. The samples were then clarified by subsequent centrifugation steps as follows: (i) 500x *g* for 5 min at 4 °C, (ii) 1000x *g* for 13 min at 4 °C, and (iii) 100.000x *g* for 30 min at 4 °C. The final supernatant was concentrated using 30kD spin filters (Merck Amicon Ultra -0.5 mL, Centrifugal filters, UFC*503096*) to a final concentration of 10 µg/µL, as judged by OD280, and subjected to SEC as indicate below. Experiments were performed in duplicates for each age group and conducted in different days.

2.2. *Size exclusion chromatography (SEC)*

SEC was performed using an ÄKTA avant system equipped with UV detection at 280 nm wavelength. The column was a Yarra-SEC-4000 column (300 x 7.8 mm, pore size 500 Å, particle size 3µm) with a SecurityGard™ cartridge GFC4000 4x3.0 mm ID as a guard column. Running conditions were temperature 4°C, flow rate 0.5 mL/min, run time of 40 min and mobile phase was 50 mM HEPES, pH 6.8, 1 mM $MgCl_2$, 1 mM DTT, 150 mM NaCl, 5 mM ATP. A standard sample (Phenomenex, ALO-3042) was injected prior to each sample to verify column performance. Sample amounts of 100 µL of a 10 mg/mL lysate were injected, corresponding to 1 mg protein extract on column. Fractions (200 µl each) were collected along with the LC separation directly in SDS buffer, to a final concentration of 4%. Thirty-nine fractions were further processed for LC-MS/MS analysis (section 4.2 for sample preparation and section 4.10 for data acquisition).

3. *In vivo proteasome inhibition*

Adult animals (12 – 14 weeks post-hatching, wph) were subjected twice to pharmacological intervention via intraperitoneal injections (IP) during a four-day period treatment. On the first and third day of the experiment (t=0 and t=48h), fish were anesthetized with 200 mg/L buffered MS-222 (PharmaQ) and gently manipulated to deliver IP of either different drugs at 500 µM or vehicle (DMSO) at a dosage of 10 µl/g body weight. After the fourth day of treatment (t=96h), fish were euthanized as previously described (section 1.1), brains harvested and used either for proteasome activity assay (section 6.2) or sample preparation for mass spectrometry (section 4.1). Different compounds were tested (see list below) in order to reach an optimal reduction of proteasome activity similar to the levels observed in old animals.

List of proteasome inhibitors tested:

| **Product** | **Company** | **Cat#** |
|---|---|---|
| Bortezomib | Sigma Aldrich | 5043140001 |
| Carfilzomib | Selleck Chemicals | S2853 |
| MG-132 | Sigma Aldrich | 474787 |
| Marizomib | Sigma Aldrich | SML1916 |

4. *Sample preparation for mass spectrometry analysis and data acquisition*

4.1. *Sample preparation for proteome analysis (N. furzeri)*

Individual brains from the fish were collected and snap frozen in liquid nitrogen (Table S1). On preparation for MS, protein amount was estimated based on fresh tissue weight (assuming 5% of protein w/w) and lysis buffer (4% SDS, 100 mM HEPES, pH8, 1mM EDTA, 100 mM DTT) was added accordingly to a final concentration of 1 µg/ µL. Samples were then vortexed (5 times) prior to sonication (Bioruptor Plus) for 10 cycles (30 sec ON/60 sec OFF) at high setting, at 4 °C. The samples were then centrifuged at 3000x *g* for 5 min at room temperature, and the supernatant transferred to 2 mL Eppendorf tubes. Reduction (15 min, 45 °C) was followed by alkylation with 20 mM iodoacetamide (IAA) for 30 min at room temperature in the dark. Protein amounts were confirmed, following an SDS-PAGE gel of 4% of each sample against an in-house cell lysate of known quantity. Between 200 and 300 µg of each sample was taken along for digestion. Proteins were precipitated overnight at -20 °C after addition of a 4x volume of ice-cold acetone. The following day, the samples were centrifuged at 20800x *g* for 30 min at 4 °C and the supernatant carefully removed. Pellets were washed twice with 1 mL ice-cold 80% (v/v) acetone in water then centrifuged at 20800x *g* at 4 °C. They were then allowed to air-dry before addition of 120 µL of digestion buffer (3M Urea, 100 mM HEPES, pH8). Samples were resuspended with sonication (as above), LysC (Wako) was added at 1:100 (w/w) enzyme:protein and digestion proceeded for 4 h at 37 °C with shaking (1000 rpm for 1 h, then 650 rpm). Samples were then diluted 1:1 with MilliQ water and trypsin (Promega) added at the same enzyme to protein ratio. Samples were further digested overnight at 37 °C with shaking (650 rpm). The following day, digests were acidified by the addition of TFA to a final concentration of 2% (v/v) and then desalted with Waters Oasis® HLB µElution Plate 30 µm (Waters Corporation, Milford, MA, USA) in the presence of a slow vacuum. In this process, the columns were conditioned with 3x100 µL solvent B (80% (v/v) acetonitrile; 0.05% (v/v) formic acid) and equilibrated with 3x100 µL solvent A (0.05% (v/v) formic acid in Milli-Q water). The samples were loaded, washed 3 times with 100 µL solvent A, and then eluted into 0.2 mL PCR tubes with 50 µL solvent B. The eluates were dried down with the speed vacuum centrifuge and dissolved at a concentration of 1 µg/µL in reconstitution buffer (5% (v/v) acetonitrile, 0.1% (v/v) formic acid in Milli-Q water). Reconstituted peptides were either analyzed directly (label-free analysis, section 4.5, used for TMT labeling, 4.6, or data independent acquisition (DIA), section 4.9).

##### 4.2. *Sample preparation for SEC fractions*

Additional DTT (to a final concentration of 50mM) in 100 mM HEPES, pH 8 was added to each fraction, followed by sonication (Bioruptor Plus) for 10 cycles (30 sec ON/60 sec OFF) at high setting, at 20 °C. Reduction and alkylation were performed as previously described (section 4.1). Protein amounts were estimated following an SDS-PAGE gel of 4% of each sample against an in-house cell lysate of known quantity. Between 10 to 40 µg of each fraction were taken along for digestion. Proteins were digested and peptide desalted as described in section 4.1 and analyzed as described in section 4.10.

##### 4.3. *Isolation of SDS insoluble protein aggregates (M. musculus)*

Individual brains from young (5 mo) and old animals (21 or 26 mo, Table S2) (between 481 and 533 mg wet tissue weight) were lysed at a protein concentration of 30 µg/ µL in lysis buffer (4% SDS, 100 mM HEPES, pH 8, 1 mM EDTA, 100 mM DTT). Samples were then vortexed (5 times) prior to sonication with a Bioruptor Plus (high setting, 20 °C, 20 cycles of 60 sec ON/30 sec OFF). Samples were then centrifuged at 20000x *g* for 5 min at room temperature, and the supernatant transferred to fresh 2 mL Eppendorf tubes (total homogenate; TH). In order to obtain pellets of aggregates, 200 µL of brain lysates (TH) were transferred to polypropylene thick wall tubes (Beckman Coulter), in duplicate, and submitted to ultracentrifugation at 100000x *g* for 30 min at 20 °C. Supernatant was transferred to a fresh tube (supernatant; SUP) and remaining pellet washed twice, by resuspension with 200 µL of lysis buffer, followed by ultracentrifugation. Supernatant from each of the washes were transferred to fresh tubes (wash 1; W1 and wash 2; W2). In order to facilitate their solubilization, pellets (SDS insoluble proteins) were then submitted to 50 µL of neat formic acid for 1 h at 37 °C with shaking (400 rpm). After incubation, samples were speed vacuum centrifuged at 45°C, resuspended in 50 µL of lysis buffer and boiled at 95°C for 10 minutes. Samples were then transferred to 0.5 mL Eppendorf tubes and rebuffered with 5 M NaOH. On preparation for MS, total homogenate (10 µg) and equal volumes of resuspended pellets (estimated amount of protein between 5 and 15 µg) for each sample were submitted to protein precipitation, digestion and clean up as described in sections 4.1.

##### 4.4. *Quantification of SDS insoluble aggregates from young and old brains*

Resuspended pellets were loaded in SDS-PAGE gel and their protein content compared. In this process, equal volumes of resuspended/rebuffered pellets were mixed with 2x loading buffer (1.5M Tris pH 6.8, 20% SDS, 85% Glycerin)), loaded in precast protein gel (BioRad, Mini-PROTEAN TGX 4-20%, 10-well) and run under constant mode (100V, 1:30h) in 1% SDS running buffer. The gel was then stained with Coomassie overnight at room temperature, with shaking, followed by extensive washing with MilliQ water. The image was acquired using the ChemiDoc XRS+ system (Bio-Rad) with the standard colorimetric settings (Image Lab 5.2.1). A high resolution image was then exported for further densitometry analysis of full length lanes with the open source software ImageJ 1.52a on a Windows 7 Professional 64-bit install (NIH) (Schneider et al., 2012). Prior to analysis, the image was converted to grey-scale 8-bit mode. In brief, a rectangular selection of same size was drawn across the full lane of each sample. Profile plots with the relative density of the contents from each rectangle were generated (function *Plot Lanes*). The generated area under the curve for each sample was measured (function *Wand*). In order to test for significant differences between young and old brains, density area values were tested for normal distribution (Shapiro-Wilk test). An unpaired parametric t-test was performed. Statistical analysis was done using built-in functions of Graphpad Prism 8.

##### 4.5. *Data acquisition for label-free analysis*

Peptides were separated using the nanoAcquity UPLC system (Waters) fitted with a trapping (nanoAcquity Symmetry $C_{18}$, 5µm, 180 µm x 20 mm) and an analytical column (nanoAcquity BEH $C_{18}$, 1.7µm, 75µm x 250mm). The outlet of the analytical column was coupled directly to an Orbitrap Fusion Lumos (Thermo Fisher Scientific) using the Proxeon nanospray source. Solvent A was water, 0.1 % (v/v) formic acid and solvent B was acetonitrile, 0.1 % (v/v) formic acid. The samples (500 ng) were loaded with a constant flow of solvent A at 5 µL/ min onto the trapping column. Trapping time was 6 minutes. Peptides were eluted via the analytical column with a constant flow of 0.3 µL/ min. During the elution step, the percentage of solvent B increased in a linear fashion from 3 % to 25 % in 30 minutes, then increased to 32 % in 5 more minutes and finally to 50 % in a further 0.1 minutes. Total runtime was 60 minutes. The peptides were introduced into the mass spectrometer via a Pico-Tip Emitter 360 µm OD x 20 µm ID; 10 µm tip (New Objective) and a spray voltage of 2.2 kV was applied. The capillary temperature was set at 300 °C. The RF lens was set to 30%. Full scan MS spectra with mass range 375-1500 *m/z* were acquired in profile mode in the Orbitrap with resolution of 120000 FWHM. The filling time was set at maximum of 50 ms with limitation of $2 \times 10^5$ ions. The "Top Speed" method was employed to take the maximum number of precursor ions (with an intensity threshold of $5 \times 10^3$) from the full scan MS for fragmentation (using HCD collision energy, 30%) and quadrupole isolation (1.4 Da window) and measurement in the ion trap, with a cycle time of 3 seconds. The MIPS (monoisotopic precursor selection) peptide algorithm was employed but with relaxed restrictions when too few precursors meeting the criteria were found. The fragmentation was performed after accumulation of $2 \times 10^3$ ions or after filling time of 300 ms for each precursor ion (whichever occurred first). MS/MS data were acquired in centroid mode, with the Rapid scan rate and a fixed first mass of 120 *m/z*. Only multiply charged ($2^+$ - $7^+$) precursor ions were selected for MS/MS. Dynamic exclusion was employed with maximum retention period of 60s and relative mass window of 10 ppm. Isotopes were excluded. Additionally, only 1 data dependent scan was performed per precursor (only the most intense charge state selected). Ions were injected for all available parallelizable time. In order to improve the mass accuracy, a lock mass correction using a background ion (*m/z* 445.12003) was applied. For data acquisition and processing of the raw data, Xcalibur 4.0 (Thermo Scientific) and Tune version 2.1 were employed.

##### 4.6. *TMT labeling*

The resuspended peptides (at 1 µg/µL) were rebuffered to pH 8.5 using 1 M HEPES prior labelling. For the total proteome experiment, 12 wph samples were used as common reference for 5 wph and 39 wph samples (see Table S1 for labeling scheme). In this case, 30 µg and 15 µg of peptides were taken for each labelling reaction, from 12 wph and 5 wph / 39 wph samples, respectively. For TPP experiments, 10 µg of peptides from each temperature point from each replicate were labelled (see Table S1 for labeling scheme). TMT-10plex reagents (Thermo Scientific) were reconstituted in 41 µL of anhydrous DMSO. TMT labeling was performed in two steps by addition of 2x of the TMT reagent per µg of peptide (e.g, 60 µg of TMT reagent for 30 µg of peptides). First, sample amount of TMT reagent was added to samples at room temperature, with shaking at 600 rpm in a thermomixer (Eppendorf) for 30 minutes. After incubation, a second portion of TMT reagent was added and incubated for another 30 minutes. After checking labelling efficiency by MS, samples were pooled (50 µg total), desalted as described in section 4.1 and subjected to high pH fractionation prior to MS analysis.

4.7. *High pH peptide fractionation for TMT labeled samples*

Offline high pH reverse phase fractionation was performed using an Agilent 1260 Infinity HPLC System equipped with a binary pump, degasser, variable wavelength UV detector (set to 220 and 254 nm), peltier-cooled autosampler (set at 10 °C) and a fraction collector. The column was a Waters XBridge C18 column (3.5 µm, 100 x 1.0 mm, Waters) with a Gemini C18, 4 x 2.0 mm SecurityGuard (Phenomenex) cartridge as a guard column. The solvent system consisted of 20 mM ammonium formate (pH 10.0) as mobile phase (A) and 100 % acetonitrile as mobile phase (B). The separation was accomplished at a mobile phase flow rate of 0.1 mL/min using a non-linear gradient from 95 % A to 40 % B in 91 min. Forty-eight fractions were collected along with the LC separation that were subsequently pooled into 16 nonconsecutive fractions. Pooled fractions were dried in a Speed-Vac and then stored at -80°C until LC-MS/MS analysis.

4.8. *Data acquisition TMT labelled, high pH fractionated samples*

For TMT experiments, fractions were resuspended in 10 µL reconstitution buffer (5% (v/v) acetonitrile, 0.1% (v/v) TFA in water) and 3 µL were injected. Peptides were separated using the nanoAcquity UPLC system (Waters) fitted with a trapping (nanoAcquity Symmetry C18, 5 µm, 180 µm x 20 mm) and an analytical column (nanoAcquity BEH C18, 2.5 µm, 75 µm x 250 mm). The outlet of the analytical column was coupled directly to an Orbitrap Fusion Lumos (Thermo Fisher Scientific) using the Proxeon nanospray source. Solvent A was water, 0.1% (v/v) formic acid and solvent B was acetonitrile, 0.1% (v/v) formic acid. The samples were loaded with a constant flow of solvent A at 5 µL/ min, onto the trapping column. Trapping time was 6 min. Peptides were eluted via the analytical column at a constant flow of 0.3 µL/ min, at 40 °C. During the elution step, the percentage of solvent B increased in a linear fashion from 5% to 7% in 10 min, then from 7 % B to 30 % B in a further 105 min and to 45 % B by 130 min. The peptides were introduced into the mass spectrometer via a Pico-Tip Emitter 360 µm OD x 20 µm ID; 10 µm tip (New Objective) and a spray voltage of 2.2 kV was applied. The capillary temperature was set at 300 °C. Full scan MS spectra with mass range 375-1500 *m/z* were acquired in profile mode in the Orbitrap with resolution of 60000 FWHM using the quadrupole isolation. The RF on the ion funnel was set to 40 %. The filling time was set at maximum of 100 ms with an AGC target of 4 x $10^5$ ions and 1 microscan. The peptide monoisotopic precursor selection (MIPS) was enabled along with relaxed restrictions if too few precursors were found. The most intense ions (instrument operated for a 3 second cycle time) from the full scan MS were selected for MS2, using quadrupole isolation and a window of 1 Da. HCD was performed with collision energy of 35%. A maximum fill time of 50 ms for each precursor ion was set. MS2 data were acquired with fixed first mass of 120 *m/z* in the ion trap. The dynamic exclusion list was with a maximum retention period of 60 sec and relative mass window of 10 ppm. The instrument was not set to inject ions for all available parallelizable time. For the MS3, the precursor selection window was set to the range 400-2000 *m/z,* with an exclude width of 18 *m/z* (high) and 5 *m/z* (low). The most intense fragments from the MS2 experiment were co-isolated (using Synchronus Precursor Selection = 8) and fragmented using HCD (65%). MS3 spectra were acquired in the Orbitrap over the mass range 100-1000 *m/z* and resolution set to 30000 FWHM. The maximum injection time was set to 105 ms and the instrument was set not to inject ions for all available parallelizable time.

4.9. *Data independent acquisition (DIA) for in vivo proteasome inhibition*

Reconstituted peptides were spiked with retention time iRT kit (Biognosys AG, Schlieren, Switzerland). Peptides were separated using the nanoAcquity UPLC system (Waters) with a trapping (nanoAcquity Symmetry C18, 5µm, 180 µm x 20 mm) and an analytical column (nanoAcquity BEH C18, 1.7µm, 75µm x 250mm). The outlet of the column was coupled to a Q exactive HF-X (Thermo Fisher Scientific) using the Proxeon nanospray source. Solvent A was water, 0.1 % FA and solvent B was acetonitrile, 0.1 % FA. Samples were loaded at constant flow of solvent A at 5 µL/min onto the trap for 6 mins. Peptides were eluted via the analytical column at 0.3 µL/min and introduced via a Pico-Tip Emitter 360 µm OD x 20 µm ID; 10 µm tip (New Objective). A spray voltage of 2.2 kV was used. During the elution step, the percentage of solvent B increased in a non-linear fashion from 0 % to 40 % in 120 minutes. Total run time was 145 minutes. The capillary temperature was set at 300 °C. The RF lens was set to 40%. MS conditions were: Full scan MS spectra with mass range 350-1650 *m/z* were acquired in profile mode in the Orbitrap with resolution of 120000 FWHM. The filling time was set at maximum of 60 ms with limitation of 3 x $10^6$ ions. DIA scans were acquired with 40 mass window segments of differing widths across the MS1 mass range. The default charge state was set to 3+. HCD fragmentation (stepped normalized collision energy; 25.5, 27, 30%) was applied and MS/MS spectra were acquired with a resolution of 30000 FWHM with a fixed first mass of 200 *m/z* after accumulation of 3 x $10^6$ ions or after filling time of 35 ms (whichever occurred first). Data were acquired in profile mode. For data acquisition and processing of the raw data Xcalibur 4.0 (Thermo Scientific) and Tune version 2.9 were employed.

4.10. *Data independent acquisition (DIA) for SEC fractions*

Data acquisition was performed as described above, including modifications in the gradient settings and MS as follows. During the elution step, the percentage of solvent B increased in a non-linear fashion from 0 % to 40 % in 60 min. Total run time was 75 min. DIA scans were acquired with 30 mass window segments. HCD fragmentation (stepped normalized collision energy; 25.5, 27, 30%) was applied and MS/MS spectra were acquired with a resolution of 30000 FWHM with a fixed first mass of 200 *m/z* after accumulation of 3 x $10^6$ ions or after filling time of 47 ms (whichever occurred first).

4.11. *Data processing for TMT labeled samples (total proteome analysis)*

TMT-10plex data were processed using Proteome Discoverer v2.0 (Thermo Fisher Scientific). Data were searched against the relevant species-specific fasta database (in-house *N. furzeri* or Uniprot database, Swissprot entry only, release 2016_01 for mouse or Uniprot database, Swissprot entry only, release 2016_01 for human) using Mascot v2.5.1 (Matrix Science) with the following settings: Enzyme was set to trypsin, with up to 1 missed cleavage. MS1 mass tolerance was set to 10 ppm and MS2 to 0.5 Da. Carbamidomethyl cysteine was set as a fixed modification and oxidation of Methionine as variable. Other modifications included the TMT-10plex modifications from the quantification method used. The quantification method was set for reporter ions quantification with HCD and MS3 (mass tolerance, 20 ppm). The false discovery rate for peptide-spectrum matches (PSMs) was set to 0.01 using Percolator (Brosch et al., 2009).

Reporter ion intensity values for the PSMs were exported and processed with procedures written in R (version 3.5.0) using R-studio (version 1.0.153), as described in (Heinze et al., 2018). Briefly, PSMs mapping to reverse or contaminant hits, or having a Mascot score below 15, or having reporter ion intensities below 1 x $10^3$ in all the relevant TMT channels were discarded. TMT channels intensities from the retained PSMs were then $\log_2$ transformed, normalised and summarised into protein group quantities by taking the median value. At least two unique peptides per protein were required for the identification and only those peptides with no missing values across all 10 channels were considered for quantification. Protein differential expression was evaluated using the limma package (Ritchie et al., 2015). Differences in protein abundances were statistically determined using the Student's t test moderated by the empirical Bayes method. P values were adjusted for multiple testing using the Benjamini-Hochberg method (FDR, denoted as "adj. p") (Benjamini and Hochberg, 1995). The results are reported in Table S3.

To obtain iBAQ values (Schwanhäusser et al., 2011) from TMT data, all samples were re-analyzed with MaxQuant 1.5.3.28. The parameters were set identically to the analyses above. For each identified peptide in each separate LC-MS analysis file, we extracted both precursor intensities as well as corresponding PSMs from the evidence files. For further analysis, we only considered PSMs that were common to both the TMT quantification approach as described above and the MaxQuant analysis, and were filtered as before. For each resulting peptide per LC-MS analysis, we calculated a peptide ratio corresponding to the median of the ratios derived from its PSMs. Given that the total area (MS1/precursor intensity) of a peptide species represents the sum of the 10 TMT channels, splitting the total area into individual channels using the TMT ratios gives the intensity portions for each channel. After splitting, we corrected for potential sampling aberrations by multiplying the area intensities per channel with the median ratio determined from the TMT ratios. For each channel and peptide, we calculated label-free scores by dividing through the number of potentially observable unique tryptic peptides per protein (criteria: peptide length 8-25 amino acids, no missed cleavage allowed). The resulting iBAQ scores of unique peptides were summed up per protein, and protein scores were normalized across samples using median normalization.

##### 4.12. *Data processing for label-free quantification (protein aggregates)*

Software MaxQuant (version 1.5.3.28) was used to search the data. The data were searched against a species-specific (*N. furzeri* in-house or Uniprot database, Swissprot entry only, release 2016_01 for mouse) database with a list of common contaminants appended. The data were searched with the following modifications: Carbamidomethyl (C) (fixed) and Oxidation (M) and Acetyl (Protein N-term) (variable). The mass error tolerance for the full scan MS spectra was set at 20 ppm and for the MS/MS spectra at 0.5 Da. A maximum of 2 missed cleavages were allowed. For aggregate analysis, iBAQ values (Schwanhäusser et al., 2011) from the MaxQuant output were used to perform quantitative analyses. Only protein groups quantified in at least two replicates per sample group were retained. To reduce technical variation, data were $\log_2$ transformed and quantile-normalized using the preprocessCore library. Differential protein expression was assessed using the limma package, as described in section 4.11. The results are reported in Table S8.

##### 4.13. *Data processing for DIA*

For experiment specific library creation, the DIA data were searched against either a *N. furzeri* in-house database (59,154 entries) or *N. furzeri* uniprot database (35,275 entries) and a list of common contaminants using Pulsar engine in Spectronaut Professional+ (version 12.0.20491.0.21234, Biognosys AG, Schlieren, Switzerland). The following modifications were included in the search: Carbamidomethyl (C) (Fixed) and Oxidation (M)/ Acetyl (Protein N-term) (Variable). A maximum of 2 missed cleavages for trypsin were allowed. The identifications were filtered to satisfy FDR of 1 % on peptide and protein level.

For the SEC experiment (section 2), a library containing all samples (young and old, in duplicates) was generated and contained 117,755 precursors. The same library was used to search data from different experiments separately. Precursor matching, protein inference, and quantification were performed in Spectronaut using default settings.

For the *in vivo* proteasome inhibition experiment (section 3), the library generated from all DIA data (DMSO and Bortezomib) contained 70,301 precursors, corresponding to 6.636 protein groups.

The protein quantity report was then exported and further data analyses and visualization were performed with R using in-house pipelines and scripts. For *in vivo* proteasome inhibition experiment, differential protein expression was assessed using the limma package, as described in section 4.11. SEC analysis was performed as described in section 8.4.

#### 5. *Sample preparation and data processing for RNA sequencing*

##### 5.1. *RNA isolation*

RNA from each sample was extracted from protein lysates using Qiazol lysis reagent (Qiagen). In brief, 1 mL of Qiazol reagent was added to 100 µL of lysate, followed by the addition of 200 µL of chloroform. Samples were mixed vigorously and centrifuged at 12000x *g* for 20 min at 4 °C, after 3 min incubation at room temperature. The upper aqueous phase was carefully transferred into a fresh tube and mixed with 1 volume of isopropyl alcohol, 0.16 volumes of Sodium acetate (2 M; pH 4.0) and 1 µL of GlycoBlue (Invitrogen™) in order to precipitate RNA. After 10 min incubation at room temperature, samples were centrifuged at 12000x *g* for 30 min at 4 °C. The supernatant was completely removed and RNA pellets were washed by adding 80% (v/v) ethanol and centrifuging at 7500x *g* for 5 min at 4 °C. The washing steps were performed twice. The resulting pellets were air-dried for no more than 5 min and dissolved in 10 µL nuclease-free water. To ensure full dissolution of RNA in water, samples were then incubated at 65 °C for 5 min, before storage at -80 °C.

##### 5.2. *Library preparation*

Sequencing of RNA samples was done using Illumina's next-generation sequencing methodology (Bentley et al., 2008). In detail, quality check and quantification of total RNA was done using the Agilent Bioanalyzer 2100 in combination with the RNA 6000 pico kit (Agilent Technologies). Total RNA library preparation was done introducing 250 ng total RNA into the Illumina's TruSeq Stranded Total RNA Library Prep Kit/ RiboZero Gold kit, following the manufacturer's instructions. Small RNA library preparation was done using Illumina's TruSeq small RNA library preparation kit following the manufacturer's description. Quality and quantity of all libraries were checked using Agilent's Bioanalyer 2100 and DNA 7500 kits.

##### 5.3. *Sequencing*

All libraries were sequenced on a HiSeq2500 running in 51 cycle/single-end/high-output mode (sequencing chemistry v3). Total RNA libraries were pooled and sequenced in three lanes. Small RNA libraries were pooled and sequenced in one lane. Sequence information was extracted in FastQ format using Illumina's bcl2fastq v.1.8.4. Sequencing of total RNA libraries resulted in around 42 mio reads per sample; sequencing of small RNA libraries in around 12 mio reads per sample.

##### 5.4. *Data processing for small RNAs*

Sequence information was extracted in FastQ format using Illumina's bcl2fastq software v1.8.3. The processing and annotation of small RNA-Seq raw data was performed using the R programming language (version 3.0.2) and the ShortRead Bioconductor package (Morgan et al., 2009). First, raw data were pre-processed with the following parameters: Quality filtering, eliminating all reads containing an "N"; Adapter trimming, by use of the function trimLRPatterns(), allowing up to 2 mismatches and using as adapter sequence "TGGAATTCTCGGGTGCCAAGGAACTCCAGTCAC". Size filtering removed all the reads with lengths shorter than 18 and longer than 33 nucleotides. Reads were aligned using Bowtie 1.1.2 (Langmead et al., 2009), resulting in a direct annotation and quantification. The alignment was divided in two steps, to allow the recognition and the annotation of the reads exceeding reference length. First, we performed alignment against the reference (mature miRNAs, *N. furzeri* reference catalogue (Baumgart et al., 2017)) with up to 2 mismatches. In this step, the reference used was the mature sequence of microRNAs. Each read was aligned using these criteria with Bowtie (settings: "-q", "--threads 8 --best", "—norc"). The remaining reads, which could not be aligned in the previous step, were used as reference for a second alignment step with Bowtie 1.1.2 (settings: -f", "-a", "--threads 8 --norc"). In this case, the annotated mature microRNAs were aligned against the reads. The information obtained in the two alignment phases was conveyed in one single table, containing a list of all the retrieved sequences and their relative counts.

RNAseq data were then processed as follow: sequences were mapped using Tophat2 (-T -x 1) (Kim et al., 2013) to the Nfu_20150522 genome. Counting was performed using featureCounts -s 0 on the genebuild_v1.150922 *N.furzeri* annotation. For both small and coding RNAs, the raw counts were analysed with DESeq2 package (Love et al., 2014) for differential expression. Differential expression was performed independently for the two comparisons shown in the work (12 wph vs. 5 wph and 39 wph vs. 12 wph) with $\alpha=0.05$. The analysis was applied to both coding and non-coding RNAs (miRNAseq data). For miRNA analysis only miRNAs up regulated with aging ($\log_2$ fold change > 0 and adj. $p < 0.05$) were considered. The results are reported in Table S4.

##### 5.5. *Longitudinal study*

*N. furzeri* longitudinal data were obtaine from: (i) an already published data set (GSE66712, 90 longitudinal fin-clip dataset from 45 animals), and (ii) newly sequenced 228 fin-clip samples from additional 114 animals of the same cohort. Sample preparation, sequencing and data analysis was done as described (Baumgart et al., 2014, Baumgart et al., 2016).

##### 5.6. *Ribosome footprinting*

RNA from additional samples (young, 6 wph; old, 26 wph; 4 replicates each age-group) were extracted and sequenced as previously described (section 5.1 and 5.3). Libraries were prepared following manufacturer's instructions without depletion of ribosomal RNA (ARTseq, Epicentre).

#### 6. *Measurements of proteasome activity*

##### 6.1. *Native gel electrophoresis and in-gel proteasome assay*

Brains from young, adult and old fish were lysed by sonication using a Bioruptor for 5 cycles (30 sec ON/60 sec OFF) at high setting, at 4 °C, in proteasome activity lysis buffer (50 mM Tris-HCl, pH 7.4, 5 mM $MgCl_2$, 5 mM ATP, 1 mM DTT, 10% glycerol). Cell lysates were then centrifuged at 16000x *g* for 10 mins, at 4°C. 40 µg of total cell lysates were subjected to native gel electrophoresis to reveal the various proteasome complexes (30S, 26S, 20S). The gel was then incubated in 50 mM Tris, pH 7.4, 5 mM $MgCl_2$, 1 mM ATP and 300 µM proteasome substrate (Suc-LLVY-AMC; UBPBio) for 30 min at 37 °C to assay chymotrypsin-like (CT-L) activity. Proteasome bands were visualized under UV. Following the CT-L activity assay, proteins in native gels were transferred to nitrocellulose membranes for immunoblotting with a monoclonal antibody against the α1, 2, 3, 5, 6 & 7 members of the 20S proteasome subunit (MCP231, Enzo BML-PW8195, 1:1000 dilution in TBS (0.1% Tween) and 5% milk). 20 μg of the same total cell lysates were subjected to SDS-PAGE, and ponceau staining was used as a loading control.

6.2. *Proteasome Peptidase Assay*

Brains from young, adult and old animals were sonicated as described in 6.1. CT-L, T-L and PGPH proteasome activities were assayed with the hydrolysis of specific fluorogenic peptides (UBPBio), namely Suc-LLVY-AMC (for CT-L activity), Boc-LRR-AMC (for T-L activity) and Z-LLE-AMC (for PGPH activity), respectively, for 1 h at 37 °C. 10 μg of total cell lysates were incubated in 50 mM Tris-HCl, pH 7.4, 5 mM MgCl2, 1 mM ATP, 1 mM DTT, 10% glycerol and 10 μM proteasome substrate for 1 h at 37 °C. Specific proteasome activity was determined as the difference between the total activity of protein extracts and the remaining activity in the presence of 20 μM MG132 (Enzo Life Sciences). Fluorescence was measured by multiple reads for 60 min at 37 °C by TECAN Kinetic Analysis (excitation 380 nm, emission 460 nm, read interval 5 min) on a Safire II microplate reader (TECAN).

*7. Immunofluorescence*

The whole brains of young (7-10 wph) and old (27-30 wph) fish were fixed overnight using a solution of paraformaldehyde (PFA) 4% in Phosphate Buffer (PB) 100 mM and cryoprotected with a solution of Sucrose 30% for 24 h. Finally, tissues were included in OCT embedding medium (Tissue-tek, Sakura) and stored at -20 °C until use. Brain sections (20 μm) were cut using a Leica cryostat, collected on Superfrost Plus slides (Menzel-Glaeser) and dried at 37 °C for 2 h, then washed in PBS (3 washings for 5 min each) to remove the embedding medium followed by an acid antigen retrieval treatment (10 mM Tri-sodium citrate, 0.05 % Tween, pH 6). The solution was brought to the boiling point in a microwave and the slides were dipped in the hot solution for 1 min, three times.

To visualize protein aggregates (Figure 3E and S4E-H), an aggresome fluorescent staining kit that binds to the beta-sheets of protein aggregates was applied. Following the protocol described from Shen and colleagues (Shen et al., 2011), a solution 1:2000 of aggresome dye in PBS was used for 3 min, followed by washing in PBS (3 washings for 5 min each), and the sections were immersed in a solution of 1% acetic acid for 30 min at room temperature to remove the excess of staining (destaining step). This step was followed by an immunofluorescence procedure: blocking solution (5% (w/v) BSA, 0.3% (v/v) Triton-X in PBS) was applied for 2 h at room temperature (RT), then the primary antibodies at the specific dilution in a solution of 1% (w/v) BSA, 0.1% (v/v) Triton in PBS, and the samples left overnight at 4°C. The following day, sections were washed in PBS (3 washings for 5 min each) to eliminate the primary antibodies solutions, and the secondary antibody was applied (Alexa-fluor 488, working dilution 1:400) for 2 hours at RT. Samples were then washed with PBS (3 washings for 5 min each) and mounted with a fluorescence mounting medium supplemented with DAPI as a nuclear staining (DAPI-Fluoroshield). Images were collected at different magnifications with a Zeiss Apotome.2 microscope provided with a ZEN 2 Pro software for the image processing, and saved as TIFF format.

To detect age-dependent gliosis (Figure S2B), double staining for S100 and Glial Fibrillary Acidic Protein (GFAP) was performed, following the standard immunohistochemistry protocols already described above: the specific primary antibodies were incubated simultaneously overnight at 4 °C, followed by PBS washings and simultaneous incubation with the specific secondary antibodies for 2h at room temperature.

See the tables below for details related to antibodies and reagents used in the histological procedures described above.

List of antibodies used:

| **Name** | **Host** | **target** | **company** | **Cat#** | **Work. Dil.** |
|---|---|---|---|---|---|
| S6 Ribosomal Protein (RPS6) | Mouse | Ribosome marker (human ribosomal protein S6) | Cell Signaling | 2317 | 1:100 |
| Anti-LAMP1 | Rabbit | Lysosome marker (human LAMP1 protein) | AbCam | Ab24170 | 1:500 |
| Anti-S100 | Rabbit | Glial cells | Dako | Z0311 | 1:400 |
| Anti-Gliarl Fibrillary Acidic Protein (GFAP) | Mouse | Neuroinflammation marker | Dako | M0761 | 1:800 |
| Proteasome 20S α1, 2, 3, 5, 6 & 7 subunits monoclonal antibody (MCP231) | Mouse | 20S proteasome | Enzo | BML-PW8195 | 1:1000 |
| Alexa-Fluor 488 | Goat | Mouse IgG | Invitrogen | A11001 | 1:400 |
| Alexa-Fluor 488 | Goat | Rabbit IgG | Invitrogen | A11008 | 1:400 |

List of reagents used for immunofluorescence:

| **Product** | **company** | **Cat#** |
|---|---|---|
| Phosphate Buffered Saline (PBS) - tablets) | Sigma-Aldrich | P4417 |
| Bovine Serum Albumine (BSA) | Sigma-Aldrich | A4503 |

| | | |
|---|---|---|
| Triton-X | Sigma-Aldrich | T8787 |
| ProteoStat Aggresome Detection Kit | Enzo Life Sciences Inc., | ENZ-51035-K100 |
| Fluoroshield with DAPI | Sigma-Aldrich | F6057 |

#### 8. Data analysis

##### 8.1. *Mapping of N. furzeri genes to human orthologues*

In order to map *N. furzeri* genes to human orthologues, the *N. furzeri* proteome (Reichwald et al., 2015) was directly mapped against the *H. sapiens* Uniprot reference proteome (accessed July 2017) using BLASTp. The hit with highest alignment score was selected as the orthologue. This procedure made it possible to assign 13484 orthologues. Teleost fish underwent whole-genome duplication after their lineage separated from the one that includes mammals, thus paralogue fish genes might diverge from the correspondent mammalian orthologue (hidden orthology). In order to detect possible candidates, the *N. furzeri* proteome was aligned against the spotted gar (*Lepisosteus oculatus*) proteome, and these orthologues consequently mapped on the human proteome. Spotted gar is a fish whose lineage diverged from teleosts before their genome duplication, thus its genome could be used as a bridge to identify hidden orthologues between teleosts and mammals (Braasch et al., 2016). The procedure resulted in the assignment of further 146 (hidden) orthologues, equivalent to 1.1% of total orthologues mapped. The mapping table is reported in Table S1.

##### 8.2. *Transcriptome / proteome comparison*

For the comparison between transcriptome and proteome two independent analyses were performed, one comparing 12 wph vs. 5 wph brain samples, and the other one comparing 39 wph vs. 12 wph samples. Differentially expressed genes and proteins were obtained for these and then plotted: for the visualization of the comparison (Figure 1D and 1E) only genes differentially expressed in both the comparisons (adj. $p < 0.05$) were considered. To identify enriched KEGG pathways (Figure 1G and 1H), a metanalysis was performed on the two dataset: p values were combined using the Fisher's Method and then adjusted for multiple testing using the Benjamini-Hochberg method (Benjamini and Hochberg, 1995). For the gene enrichment, genes contained in each quadrant were analyzed separately with WebGestalt using KEGG pathways with a cut-off of $FDR < 0.05$. To obtain the correlation between protein and transcript levels for the different time points normalized counts (obtained from DESeq2 package) and IBAQ (obtained from MaxQuant) were respectively used. Pearson correlation was calculated for every sample pair (Figure 1C).

##### 8.3. *Analysis of SEC-MS data*

The same complex definitions used for stoichiometry analysis were used (Table S7). Protein quantification report exported from Spectronaut (section 4.13) was processed using R 3.5.3, using in-house generated functions. Intensity values for each protein across the 39 fractions were normalized, dividing each value by the total sum of intensities along the fractions. Only complexes with at least 5 subunits identified were considered for further analysis. For each experiment, the distribution of pairwise Pearson's correlation between members of the same protein complex was analyzed. A set of randomly defined protein complexes of same length was generated as a control for this analysis. For each complex, the median abundance of all the quantified subunits was used to generate the complex profile across fractions. All the complex profiles were scaled to the max value (set to 1) to make profiles comparable across experiments. A heatmap of the different protein complexes across the fractions were generated using the R package *pheatmap*. The results of SEC-MS data are reported in Table S6.

8.4. *Protein complex analysis*

Protein complex stoichiometries were analyzed using method and complex definitions described in (Ori et al., 2016). Briefly, protein intensities obtained from TMT experiments were assigned to the respective protein complex and normalized by the mean complex abundance (estimated using the trimmed mean protein intensity of all complex members). In order to identify stoichiometry changes, the complex-normalized protein matrix was used for differential expression analysis using limma as described in section 4.11. Only protein complexes that had at least 5 members quantified were retained for analysis. Protein complexes were considered as "affected" if they had at least two members that showed differential expression across the conditions tested (adj. $p < 0.05$ and absolute $\log_2$ fold change > 0.5), unless otherwise stated. The results are reported in Table S7 for *N. furzeri* and Table S9 for *in vivo* proteasome inhibition experiment.

8.5. *Calculation of biophysical properties for aggregates*

Predictions of biophysical properties from the amino acid sequence of proteins in the dataset were computed using the cleverSuite (Klus et al., 2014) (chaperone requirement, intrinsic disorder) and s2D (Sormanni et al., 2015) (intrinsic disorder) classifiers. A biophysical property score is then assigned to proteins from the output of the classifier. CleverSuite is trained on a dataset of proteins that show the property under study (e.g. proteins requiring chaperons to fold, positive database), versus a 'negative' database (e.g. self-folding proteins). The minimal output of the classifier is a label according to the similarity to one of the two datasets (positive, negative, or indeterminate) and the associated probability $p$ of correct classification, that can be converted to the score:

$$s_{bp}^{cS} = \begin{cases} -1 \cdot p(s_{bp} = x) \text{ if negative} \\ 0 \text{ if indeterminate} \\ p(s_{bp} = x) \text{ if positive} \end{cases} .$$

The s2D classifier determines the likelihood that a residue in a protein sequence would be included in an α-helix, β-sheet, or random coil; its minimal output is a string of predicted secondary conformations for each aminoacid of the protein. A global score of disorder could be then determined from the number of residues predicted in a random coil state as follows:

$$S_c^{s2D} = \frac{n_c}{N}$$

where $n_c$ is the number of residues predicted to assume a random coil conformation and N is the total number of residues. Analysis of correlation between predicted scores and $T_m$ or protein enrichment in aggregates was performed using custom Python 3.5 scripts. Statistical analysis was done using built-in functions of Graphpad Prism 7.

8.6. *Analysis of N. furzeri ribosome footprinting data*

Ribosome footprinting data were generated from young and aged killifish brain extracts (6 wph and 26 wph, respectively, see section 5.6). Footprints were mapped to the *N. furzeri* reference genome (NotFur1) using the STAR sequence aligner (version 2.7.1a, (Dobin et al., 2013)). The relative abundance of footprints per gene was quantified with RSEM (version 1.3.1, (Li and Dewey, 2011)).

8.7. *Cox-Hazard model for longitudinal RNAseq data*

In order to isolate predictive factors of aging, survival analysis was performed to correlate mortality risk to gene expression. Cox-Hazard Model (also called cox regression) was applied using the formula:

$$h(t|\Delta_{ij}) = h_0(t)exp(c_i\Delta_{ij})$$

$$\Delta_{ij} = log_2(g_{ij(20)}/g_{ij(10)})$$

were $h_0(t)$ is the *baseline hazard function*, $g_{ij(20)}$ is the expression of gene *i* in the sample *j* at age 20 weeks, $g_{ij(10)}$ is the expression of gene *i* in the sample *j* at age 10 weeks and $c_i$ is a coefficient.

If $c_i > 0$, mortality increase the larger $\Delta_{ij}$, if $c_i < 0$, mortality decreases the larger $\Delta_{ij}$.

The analysis was performed using the *survival* package in R environment for all 159 animals that survived longer than 20 weeks. Normalized pseudo-counts obtained from DESeq2 were used as input and the $c_i$ values were used as input for gene set enrichment using *gage* (Luo et al., 2009). The results of the Cox-Hazard analysis are reported in Table S10.

8.8. *Data visualization*

Throughout the manuscript, in boxplots the horizontal line represent the median, the bottom and top of the box the 25th and 75th percentile, respectively, and the whiskers extend 1.5 folds the interquartile range.

8.9. *Data availability*

The mass spectrometry proteomics data have been deposited to the ProteomeXchange Consortium via the PRIDE (Vizcaino et al., 2016) partner repository with the following dataset identifiers: PXD012314, PXD016459 and PXD016587.

The sequencing data discussed in this publication have been deposited in NCBI's Gene Expression Omnibus and are accessible through GEO with the following dataset identifiers: GSE52462, GSE66712, GSE125373 and GSE124638.

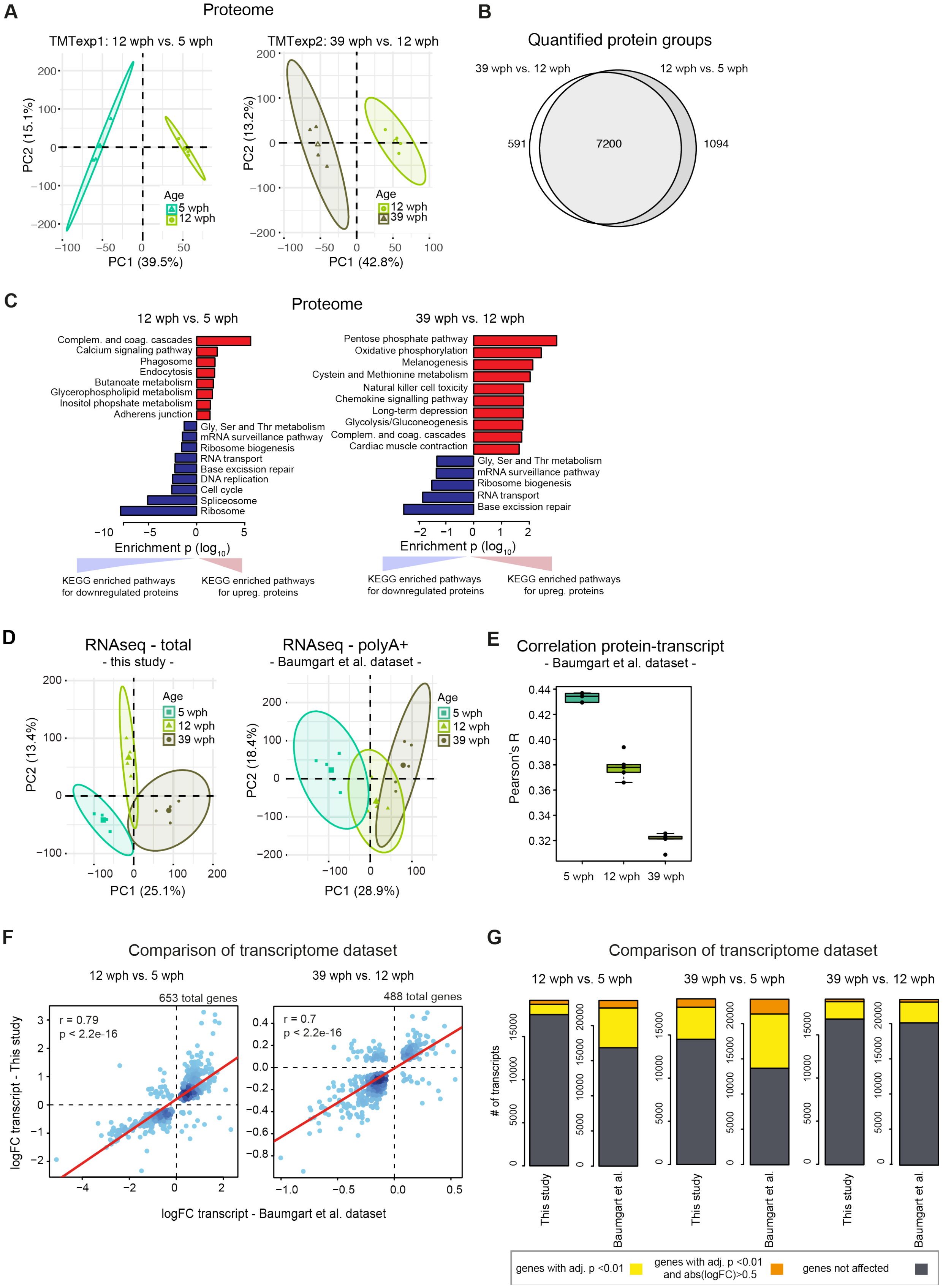
A
Proteome
TMTexp1: 12 wph vs. 5 wph
TMTexp2: 39 wph vs. 12 wph
PC2 (15.1%)
PC1 (39.5%)
PC2 (13.2%)
PC1 (42.8%)
Age
5 wph
12 wph
39 wph
B
Quantified protein groups
39 wph vs. 12 wph
12 wph vs. 5 wph
591
7200
1094
C
Proteome
12 wph vs. 5 wph
Complem. and coag. cascades
Calcium signaling pathway
Phagosome
Endocytosis
Butanoate metabolism
Glycerophospholipid metabolism
Inositol phosphate metabolism
Adherens junction
Gly, Ser and Thr metabolism
mRNA surveillance pathway
Ribosome biogenesis
RNA transport
Base excission repair
DNA replication
Cell cycle
Spliceosome
Ribosome
Enrichment p (log10)
KEGG enriched pathways for downregulated proteins
KEGG enriched pathways for upreg. proteins
39 wph vs. 12 wph
Pentose phosphate pathway
Oxidative phosphorylation
Melanogenesis
Cystein and Methionine metabolism
Natural killer cell toxicity
Chemokine signalling pathway
Long-term depression
Glycolysis/Gluconeogenesis
Complem. and coag. cascades
Cardiac muscle contraction
Gly, Ser and Thr metabolism
mRNA surveillance pathway
Ribosome biogenesis
RNA transport
Base excission repair
Enrichment p (log10)
KEGG enriched pathways for downregulated proteins
KEGG enriched pathways for upreg. proteins
D
RNAseq - total
- this study -
PC2 (13.4%)
PC1 (25.1%)
RNAseq - polyA+
- Baumgart et al. dataset -
PC2 (18.4%)
PC1 (28.9%)
Age
5 wph
12 wph
39 wph
E
Correlation protein-transcript
- Baumgart et al. dataset -
Pearson's R
5 wph
12 wph
39 wph
F
Comparison of transcriptome dataset
12 wph vs. 5 wph
653 total genes
r = 0.79
p < 2.2e-16
39 wph vs. 12 wph
488 total genes
r = 0.7
p < 2.2e-16
logFC transcript - This study
logFC transcript - Baumgart et al. dataset
G
Comparison of transcriptome dataset
12 wph vs. 5 wph
39 wph vs. 5 wph
39 wph vs. 12 wph
### of transcripts
This study
Baumgart et al.
genes with adj. p <0.01
genes with adj. p <0.01 and abs(logFC)>0.5
genes not affected

**Figure S1 related to Figure 1.**

**Proteome analysis of killifish brain aging and comparison of transcriptome dataset. (A)** Principal component analysis (PCA) of brain samples based on proteome profiles obtained by Tandem Mass Tag (TMT) quantification for the 12 wph vs. 5 wph and 39 wph vs. 12 wph comparisons. For each sample group, the four samples displaying the highest within group correlation were selected and used for differential expression analysis. The smaller dots represent individual samples and the larger dots the centroids of each age-matched group. Ellipses represent 95% confidence intervals. The percentage of variance explained by the first two PC axes is reported in the axis titles. **(B)** Overlap between the quantified protein groups across the two TMT experiments. Only proteins quantified with at least two unique (proteotypic) peptides were considered. **(C)** Barplots representing enriched KEGG pathways among proteins affected by aging in killifish brain. Pathway enrichment was performed using *gage* (Luo et al., 2009). Significant pathways enriched among up-regulated (red) or down-regulated proteins (blue) are shown ($p < 0.05$). The complete list of enriched pathway is reported in Table S3. **(D)** Principal component analysis (PCA) of brain samples based on total RNA ("this study", same sample used for proteome analysis) or polyA+ RNA sequencing ("Baumgart et al.", (Baumgart et al., 2014)). The smaller dots represent individual samples and the larger dots the centroids of each age-matched group. Ellipses represent 95% confidence intervals. The percentage of variance explained by the first two PC axes is reported in the axis titles. **(E)** Correlation between transcript and protein during aging. RPKM and iBAQ values were used to estimate transcript and protein levels from RNAseq (Baumgart et al., 2014) and TMT-based proteomics data (this study). Since in this case samples were not matched, proteomic data from each individual sample were compared against combined RPKM values obtained from the average of the samples for each group. The ANOVA test was performed to evaluate significance among the age groups (mean correlation at 5 wph: 0.43; at 12 wph: 0.38; and at 39 wph: 0.32; $p = 2.02e-12$). **(F)** Comparison between RNA-seq data obtained in this work and previous data obtained from (Baumgart et al., 2014). Fold changes ($\log_2$) were compared and plotted for both the aging comparisons (12 wph vs. 5 wph and 39 wph vs. 12 wph) for significantly affected transcripts in both comparisons (adj. $p < 0.01$). **(G)** Statistics of differentially expressed genes in the two RNA-seq datasets. The same datasets used for G were analysed. Differentially expressed genes obtained for three different age comparisons were compared between the datasets. Dark grey boxes correspond to not-affected transcripts, while yellow (adj. $p < 0.01$) and orange (adj. $p < 0.01$ and absolute $\log_2$ fold change $> 0.5$) boxes to differentially expressed genes. The complete list of quantified transcripts is reported in Table S4.

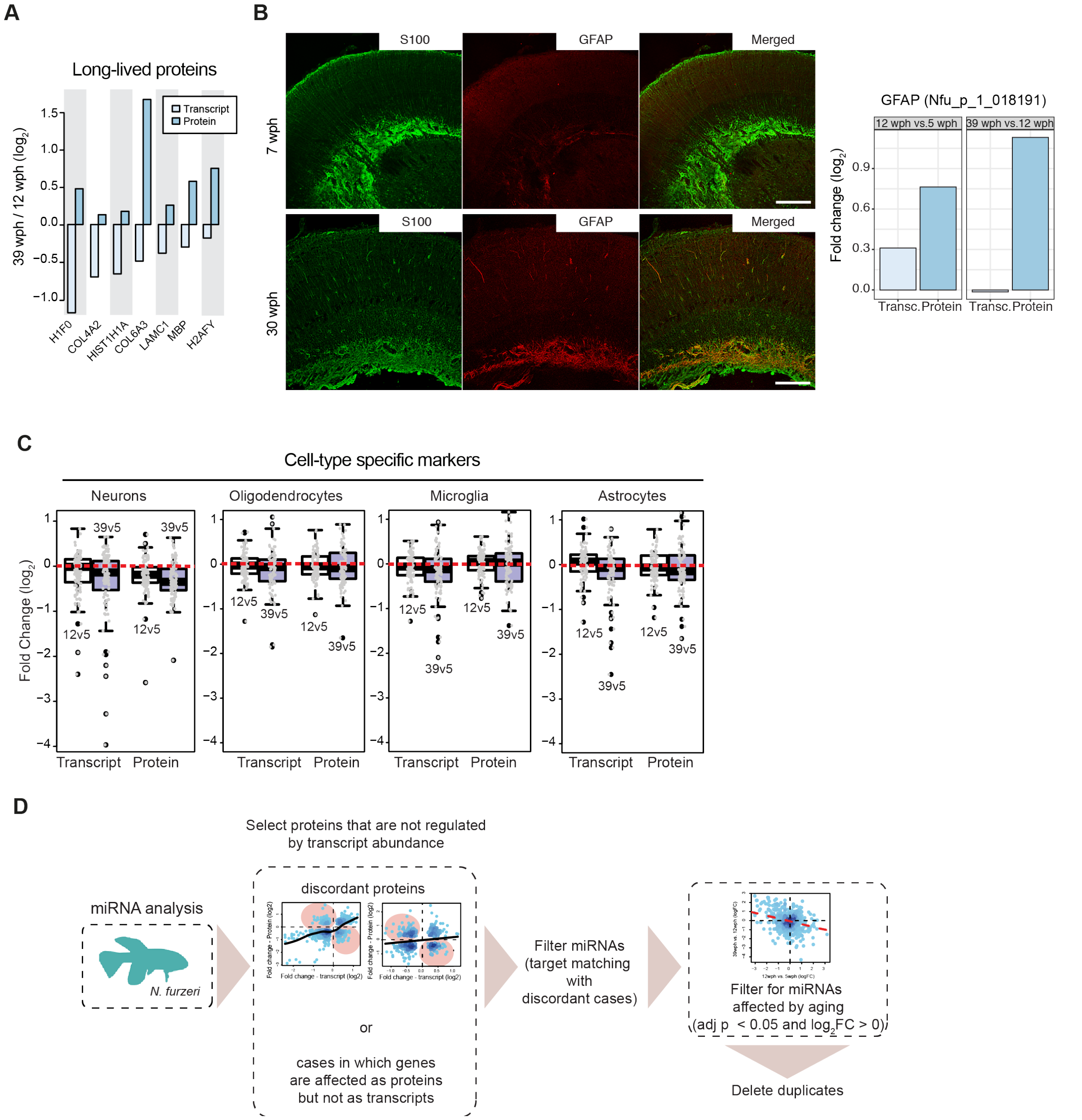


**Figure S2 related to Figure 1.**

**Validation of discordant transcript and protein changes during aging. (A)** Analysis of age-related changes of abundance for long-lived proteins. Average fold changes ($log_2$) are displayed as bars for transcripts (light blue) and proteins (dark blue), for a subset of extremely long-lived proteins identified by (Toyama et al., 2013). **(B)** Validation of an increased level of Glial Fibrillary Acidic Protein (GFAP) in old killifish brain that manifests exclusively at the protein level. Double immunostaining for S100 (green) and GFAP (red) in the central region of the optic tectum of 7 wph fish (upper panels) vs. 30 wph fish (lower panels). Scale bar = 100 µm. Average transcript and protein fold changes ($log_2$) are displayed as bars for transcript (light blue) and protein (dark blue) levels of GFAP. **(C)** Age-related changes of markers for specific cell types in the brain of *N.furzeri*. Fold changes for different age pairwise comparisons (12 wph vs. 5 wph in white; 39

wph vs. 5 wph in blue) were plotted using the cell-type markers from {Sharma, 2015 #308} (in particular neuronal, oligodendrocytes, microglia, and astrocytes markers). For each cell type, the top 100 marker genes (ranked according to adj. p value) were plotted. Both transcriptome and proteome fold changes are displayed. **(D)** Workflow for the identification of proteins potentially affected by miRNA regulation during aging. For this analysis, proteins were divided into two groups: (i) proteins affected by aging (adj. $p < 0.05$) whose abundance change could be explained by transcript expression change, and (ii) proteins affected by aging whose transcript was either regulated but with opposite fold change (discordant cases) or not regulated at all. miRNA analysis was performed only on the latter. To obtain a list of miRNAs affected by aging, differential expression was performed on both the age comparisons (12 wph vs. 5 wph and 39 wph vs. 12 wph) separately. The resulting list of miRNAs affected by aging is available in Table S4.

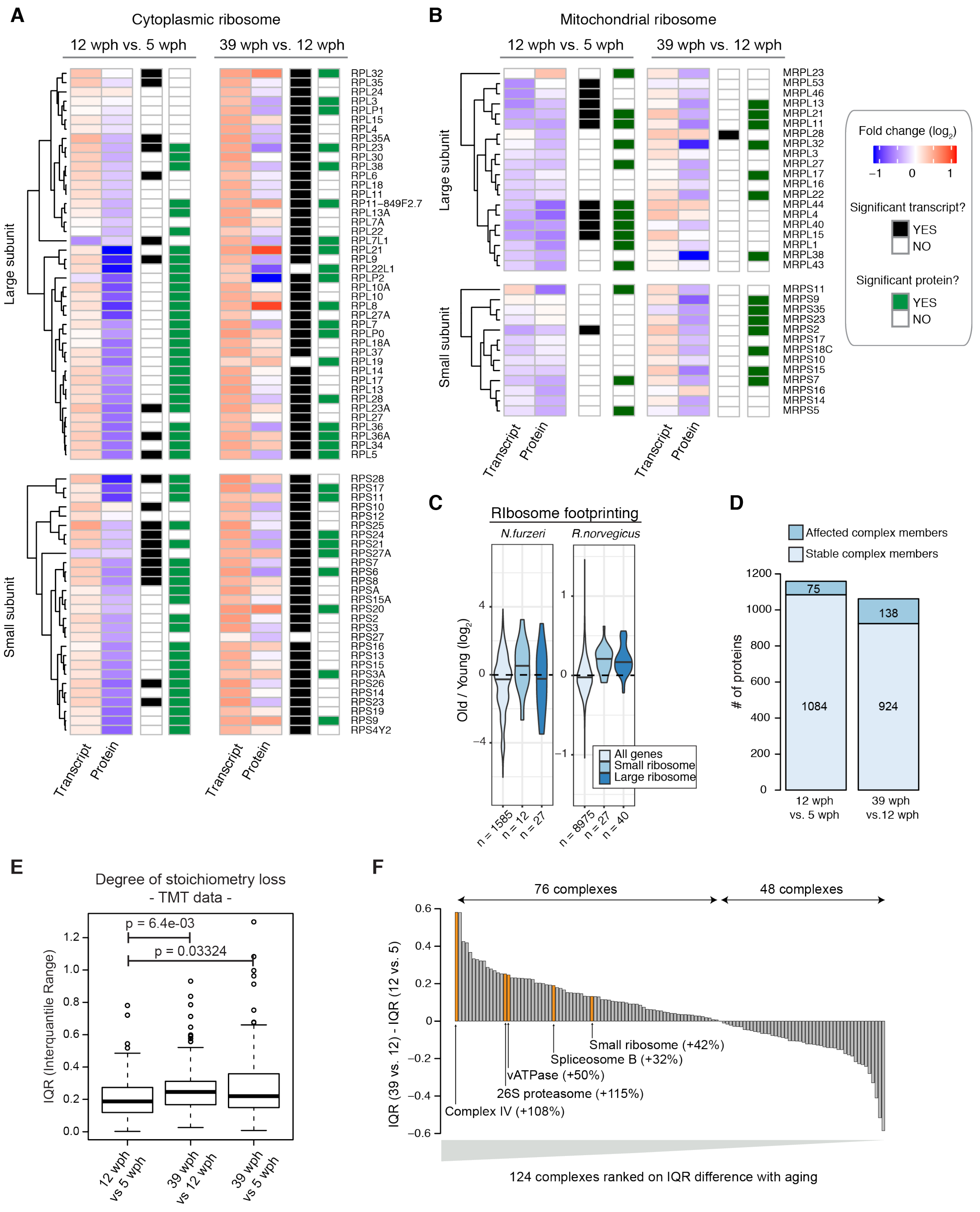


**Figure S3 related to Figure 2.**

**Detailed view of age-related abundance changes for ribosomal transcript and proteins.**

**(A and B)** Heatmap showing transcript and protein fold changes for members of the cytoplasmic (A) and mitochondrial (B) ribosome. Genes are annotated according to whether they are significantly affected at the transcript (adj. $p < 0.05$, black and white heatmap) or protein (adj. p <

0.05, black and green heatmap) level. **(C)** Ribosome footprinting analysis of young and old killifish brain. Ribosome foorprinting was performed from young (6 wph) and old (26 wph) killifish brains (n=4 per age group). Fold changes were estimated for each gene from mean TPMs of young and old samples. For comparison, age-related changes in translation output measured in rat brain from (Ori et al., 2015) are shown. **(D)** Statistics of members of protein complexes undergoing stoichiometry changes with aging. Members were considered as affected when their abundance differed significantly (adj. $p < 0.05$) from the mean of the protein complex to which they belong, as described in (Ori et al., 2016). Only protein complexes that had at least 5 members quantified were considered for each comparison. The complete list of stoichiometry changes is available in Table S7. **(E)** Degree of stoichiometry loss across different age comparisons. The inter-quantile range (IQR) of fold changes for members of the same protein complex was used to estimate the degree of stoichiometry loss (Janssens et al., 2015). All the measurements were performed in a single TMT experiment (n=3 for each age group) and p values were calculated using Wilcoxon Rank Sum test. **(F)** Bar plot showing the ranking of protein complexes based on the difference in protein level IQR for each complex between the 39 wph vs. 12 wph and 12 wph vs. 5 wph comparison. Selected complexes are highlighted and the percent of IQR increase between the two age comparisons is indicated in brackets.

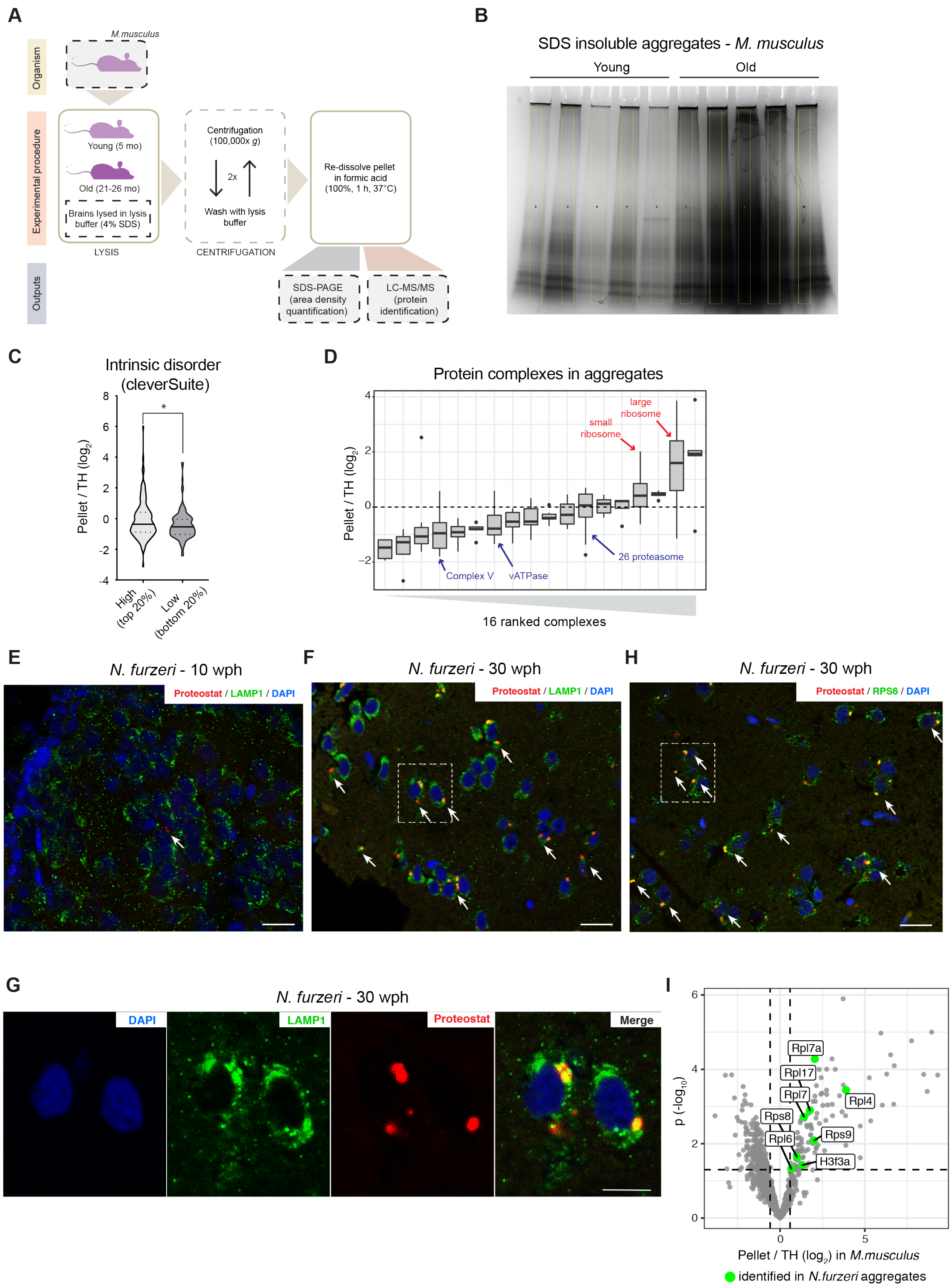


**Figure S4 related to Figure 3.**

**Biophysical properties of protein enriched in protein aggregates and validation of ribosome aggregation by immunofluorescence in old killifish brain. (A)** Workflow for the isolation of SDS insoluble aggregates from mouse brain lysates. **(B)** Coomassie stained SDS-PAGE gel used for quantification of SDS-insoluble aggregates obtained from young and old mouse brain shown in Figure 3B. For each sample, a region of interest of the same area was selected for densitometry quantification (highlighted in yellow). **(C)** Plot representing the fold enrichment in the insoluble fraction of protein with either high (top 20% scores) or low (bottom 20% scores) content of intrinsically disordered region, as predicted with the cleverSuite classifier. Violin plots: the solid line shows the median, the dotted lines the interquartile ranges. *: $p<0.05$, Kolmogorov-Smirnov test. **(D)** Boxplots of complexes ranked according to the median enrichment of protein complex members in the aggregates. Only complexes with at least 3 members quantified were considered. **(E-G)** Double-labeling of telencephalic sections of *N. furzeri* with Proteostat as a marker of aggregated proteins (red) and anti-LAMP1 as lysosomal marker (E,F and G, green, young versus old fish) and RPS6 to evidence the ribosomal component of protein aggregates (H, green, old fish). Nuclear counterstaining was performed with DAPI (blue). The region of interest shown in Figure 3E and in **G** are indicated by a white box. Scale bars = 20 µm. **(G)** Magnification of the selected area in F showing a detail of the co-localization between lysosomal structures (green) and protein aggregates (red) in the old brain. Scale bar = 10 µm. **(I)** Ribosomal proteins enriched in mouse aggregates can also be identified in *N. furzeri* aggregates. Volcano plot based on protein quantification by label-free mass spectrometry depicting the enrichment of specific proteins in protein aggregates in mice (same as shown in Figure 3D). The x-axis indicates the $\log_2$ ratio between protein abundance in aggregates (Pellet) and starting total homogenate (TH). The horizontal dashed line indicates a p value cut-off of 0.05 and vertical lines a $\log_2$ fold change cut off of ± 0.5. Proteins identified in mouse and killifish aggregates are highlighted as green dots (the entire list of proteins identified in killifish aggregates with at least 2 unique peptides in at least one replicate is reported in Table S8).

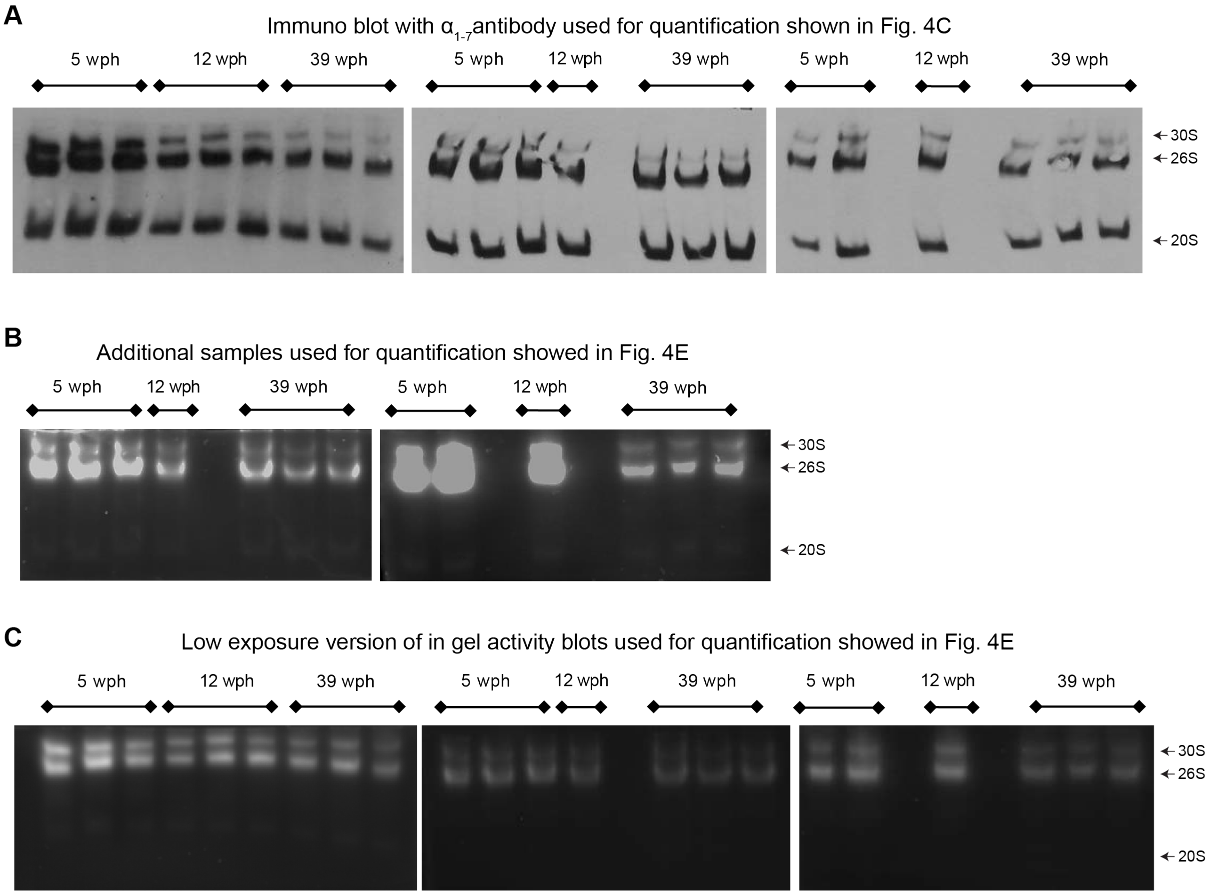


**Figure S5 related to Figure 4.**

**Decreased proteasome activity in old killifish brain. (A)** Immunoblot following native gel electrophoresis in brain extracts of killifish of different ages using an antibody recognizing the α subunits (1-7) of the proteasome. **(B)** In-gel proteasome activity following native gel electrophoresis in brain extracts of killifish of different ages. This independent group of samples was quantified together with the ones displayed in Figure 4D, and the quantification results are displayed in Figure 4E. **(C)** Low exposure of the blots show in Figures 4D and S5B. The loading of the native PAGE was based on protein quantification using the Bradford method, as described in (Myeku et al., 2011), and, additionally, confirmed by Ponceau S performed on an SDS-PAGE that was run in parallel to the native-PAGE using the same samples.

**Table S2 related to Figure 1.**

Statistics of transcriptome and proteome changes during *N. furzeri* brain aging.

| Protein groups | | | |
|---|---|---|---|
| | 12 vs. 5 stable | 12 vs. 5 up | 12 vs. 5 down |
| 39 vs. 12 stable | 3462 | 493 | 837 |
| 39 vs. 12 down | 690 | 111 | 252 |
| 39 vs. 12 up | 978 | 127 | 250 |

Out of 7200 detected proteins (overlap between the two conditions).

| Transcripts | | | |
|---|---|---|---|
| | 12 vs. 5 stable | 12 vs. 5 up | 12 vs. 5 down |
| 39 vs. 12 stable | 18557 | 605 | 244 |
| 39 vs. 12 down | 786 | 63 | 25 |
| 39 vs. 12 up | 885 | 144 | 92 |

Out of 21813 transcripts (overlap between the two conditions).

Legend

| |
|---|
| Stable = not significant |
| Down = adj. $p < 0.05$ & $\log_2$ FC < 0 |
| Up = adj. $p < 0.05$ & $\log_2$ FC > 0 |

**The following supplemental tables are provided as separate Excel files:**

Table S1: Experimental animals
Table S3: *N. furzeri* proteome aging
Table S4: *N. furzeri* transcriptome aging
Table S5: Comparison of transcriptome and proteome (*N. furzeri*)
Table S6: Size Exclusion Chromatography – mass spectrometry (*N. furzeri*)
Table S7: Protein complex analysis for *N. furzeri*
Table S8: Protein aggregates (*M. musculus* and *N. furzeri*)
Table S9: *In vivo* proteasome inhibition – proteome and protein complex analysis (*N. furzeri*)
Table S10: Longitudinal study (*N. furzeri*)

**Table S1 related to Figure 1-6.**

Experimental animals and cell line

- Tab1. List of animals (*N. furzeri*) for transcriptome and proteome analysis
- Tab2. List of animals (*M. musculus*) for protein aggregate analysis
- Tab3. List of animals (*N. furzeri*) for protein aggregate analysis
- Tab4. List of animals (*N. furzeri*) for SEC
- Tab5. List of animals (*N. furzeri*) for proteasome activity assay
- Tab6. List of animals (*N. furzeri*) for *in vivo* proteasome inhibition
- Tab7. Mapping of *N. furzeri* genes to human orthologues

**Table S3 related to Figure 1.**

*N. furzeri* proteome aging

- Tab1. Limma output for 12wph vs. 5wph comparison
- Tab2. Limma output for 39wph vs. 12wph comparison
- Tab3. KEGG pathways enriched in up regulated cases (12wph vs. 5wph)
- Tab4. KEGG pathways enriched in down regulated cases (12wph vs. 5wph)
- Tab5. KEGG pathways enriched in up regulated cases (39wph vs. 12wph)
- Tab6. KEGG pathways enriched in downregulated cases (39wph vs. 12wph)

**Table S4 related to Figure 1.**

*N. furzeri* transcriptome aging

- Tab1. DeSeq2 output for 12wph vs. 5wph comparison (total RNA dataset)
- Tab2. DeSeq2 output for 39wph vs. 12wph comparison (total RNA dataset)
- Tab3. DeSeq2 output for 12wph vs. 5wph comparison (polyA+ dataset, Baumgart *et al.*)
- Tab4. DeSeq2 output for 39wph vs. 12wph comparison (polyA+ dataset, Baumgart *et al.*)
- Tab5. miRNA

**Table S5 related to Figure 1.**

Comparison of transcriptome and proteome (*N. furzeri*)

- Tab1. Comparison transcriptome vs. proteome 12wph vs. 5wph
- Tab2. Comparison transcriptome vs. proteome 39wph vs. 12wph
- Tab3. KEGG pathways enriched for concordant cases in 12wph vs. 5wph (Up transcript / Up protein)
- Tab4. KEGG pathways enriched for discordant cases in 12wph vs. 5wph (Down transcript / Up protein)
- Tab5. KEGG pathways enriched for concordant cases in 12wph vs. 5wph (Down transcript / Down protein)
- Tab6. KEGG pathways enriched for discordant cases in 12wph vs. 5wph (Up transcript / Down protein)
- Tab7. KEGG pathways enriched for concordant cases in 39wph vs. 12wph (Up transcript / Up protein)
- Tab8. KEGG pathways enriched for discordant cases in 39wph vs. 12wph (Down transcript / Up protein)
- Tab9. KEGG pathways enriched for concordant cases in 39wph vs. 12wph (Down transcript / Down protein)

Tab10. KEGG pathways enriched for discordant cases in 39wph vs. 12wph (Up transcript / Down protein)

**Table S6 related to Figure 2.**

Size Exclusion Chromatography – mass spectrometry (*N. furzeri*)

Tab1. Tab1. Protein complex definition
Tab2. Median abundance of all the quantified subunits used to generate the complex profile across fractions

**Table S7 related to Figure 2.**

Protein complex analysis for *N. furzeri*

Tab1. Protein complex definitions
Tab2. Protein complex analysis 12wph vs. 5wph
Tab3. Protein complex analysis 39wph vs. 12wph
Tab4. Protein complex IQR between ages (Transcript - total RNA dataset)
Tab5. Protein complex IQR between ages (Proteins)

**Table S8 related to Figure 3.**

Protein aggregates (*M. musculus* and *N. furzeri*)

Tab1. Limma output for aggregates enriched in old brains (*M. musculus*) (Pellet vs. Total Homogenate)
Tab2. List of protein identified in aggregates of old brains (*N. furzeri)* with at least 2 unique peptides and in at least one replicate)

**Table S9 related to Figure 5.**

*In vivo* proteasome inhibition – proteome and protein complex analysis (*N. furzeri*)

Tab1. Protein complex definition
Tab2. Limma output for Bortezomib vs. DMSO comparison
Tab3. Protein complex analysis Bortezomib vs. DMSO comparison

**Table S10 related to Figure 6.**

Longitudinal study (*N. furzeri*)

Tab1. Cox Harzard analysis - $c_i$ coefficients
Tab2. KEGG pathways enriched for risk factors
Tab3. KEGG pathways enriched for protective factors
